# Resilience and vulnerabilities of tumor cells under purine shortage stress

**DOI:** 10.1101/2025.03.19.644180

**Authors:** Jianpeng Yu, Chen Jin, Cheng Su, David Moon, Michael Sun, Hong Zhang, Xue Jiang, Fan Zhang, Nomi Tserentsoodol, Michelle L. Bowie, Christopher J Pirozzi, Daniel J. George, Robert Wild, Xia Gao, David M Ashley, Yiping He, Jiaoti Huang

**Affiliations:** Department of Pathology, Duke University School of Medicine, Durham, North Carolina, USA; The Preston Tisch Brain Tumor Center, Duke University, Durham, North Carolina, USA; Department of Neurosurgery, Duke University School of Medicine, Durham, North Carolina, USA; Department of Medicine, Duke University School of Medicine, Durham, North Carolina, USA; Dracen Pharmaceuticals, Inc., San Diego, California, USA; Department of Molecular and Cellular Biology, Baylor College of Medicine, Houston, Texas, USA; USDA/ARS Children’s Nutrition Research Center, Department of Pediatrics, Baylor College of Medicine, Houston, Texas, USA

**Keywords:** prostate cancer, purine biosynthesis, MTAP, glioma, DRP-104

## Abstract

Purine metabolism is a promising therapeutic target in cancer; however how cancer cells respond to purine shortage,particularly their adaptation and vulnerabilities, remains unclear. Using the recently developed purine shortage-inducing prodrug DRP-104 and genetic approaches, we investigated these responses in prostate, lung and glioma cancer models. We demonstrate that when de novo purine biosynthesis is compromised, cancer cells employ microtubules to assemble purinosomes, multi-protein complexes of de novo purine biosynthesis enzymes that enhance purine biosynthesis efficiency. While this process enables tumor cells to adapt to purine shortage stress, it also renders them more susceptible to the microtubule-stabilizing chemotherapeutic drug Docetaxel. Furthermore, we show that although cancer cells primarily rely on de novo purine biosynthesis, they also exploit Methylthioadenosine Phosphorylase (MTAP)-mediated purine salvage as a crucial alternative source of purine supply, especially under purine shortage stress. In support of this finding, combining DRP-104 with an MTAP inhibitor significantly enhances tumor suppression in prostate cancer (PCa) models in vivo. Finally, despite the resilience of the purine supply machinery, purine shortage-stressed tumor cells exhibit increased DNA damage and activation of the cGAS-STING pathway, which may contribute to impaired immunoevasion and provide a molecular basis of the previously observed DRP-104-induced anti-tumor immunity. Together, these findings reveal purinosome assembly and purine salvage as key mechanisms of cancer cell adaptation and resilience to purine shortage while identifying microtubules, MTAP, and immunoevasion deficits as therapeutic vulnerabilities.

## Introduction

Glutamine is a major source of carbon for cells and also provides nitrogen for the synthesis of nucleotides and other molecules to promote cancer progression ^1–5^. Our previous work has established glutamine as a key driver of prostate cancer (PCa) and highlighted both the therapeutic potential and, more importantly, the challenges of effectively targeting its metabolism ^6,7^. DON (6-diazo-5-oxo-norleucine), a glutamine antagonist that broadly and irreversibly inhibits multiple enzymes involved in glutamine metabolism, has demonstrated broad anti-tumor activity in patients. However, its clinical utility is limited by gastrointestinal toxicity ^8^. Recently, prodrugs of DON have been developed that are inactive in normal cells but converted to functional DON in tumor cells due to the increased expression of certain enzymes in tumors ^8–10^. Specifically, one such prodrug, DRP-104 (Sirpiglenastat), has shown therapeutic efficacy in lymphoma, colorectal cancer, lung cancer, and pancreatic cancer models, and has been fast-tracked by the FDA for treating non-small cell lung cancer ^10–13^.

Advanced, hormone-sensitive PCa can be treated with hormonal therapy that inhibits androgen receptor (AR) functions. However the tumor eventually recurs as castration-resistant PCa (CRPC). Most CRPCs retain an adenocarcinoma histology and respond to newer AR antagonists such as enzalutamide, although resistance inevitably develops ^14–18^. Some CRPCs are histologically classified as small cell carcinoma or neuroendocrine (NE) PCa (namely NEPC), which are rapidly lethal due to lack of effective treatments^19^. The challenge in treating PCa is further compounded by its extreme intra-tumor heterogeneity. For instance, adenocarcinomas frequently also contain NE-like tumor cells ^20^, and even cancers in the hormone-sensitive stage harbor pre-existing CRPC-like cells that are destined for therapy resistance and tumor progression^21^. These features contribute to hormonal therapy failure and underscore the urgent need to develop novel, AR-independent therapies.

We have shown that advanced PCa is addicted to glutamine and inhibition of glutamine catabolism has the potential to be an effective, AR-independent therapeutic strategy ^6,7^. We have shown that DRP-104, a pan glutamine antagonist, inhibits PCa growth with minimal toxicity to normal tissues ^22^. While the DRP-104-derived effector metabolite, DON, is expected to disrupt at least seven glutamine-utilizing pathways by inhibiting as many as 10 enzymes ^23–32^, it remains unclear which of these pathways are most critical to tumor survival.

Addressing this question can guide the development of novel treatments and help identify patients who may benefit most from specific therapies. In this context, recent studies, including ours, have demonstrated that DRP-104 effectively disrupts purine supply in tumor cells^12,22^. These findings, along with prior observations that among DON’s pharmacological targets, the de novo purine biosynthesis enzyme FGAMS is the most sensitive to inhibition (with the lowest Ki) ^23–32^, nominate DRP-104 as a potential blocker of purine supply (i.e., an inducer of purine shortage stress). Since purines are essential building blocks and energy sources for cancer cells, purine biosynthesis, primarily de novo purine biosynthesis, has been explored for cancer treatment ^33^. Purine shortage can be induced by certain anti-cancer drugs such as Methotrexate, an inhibitor of Dihydrofolate Reductase (DHFR), although toxicity remains a major limitation ^34^. The establishment of DRP-104, a low-toxicity prodrug, as an inducer of purine shortage thus provides a new avenue for targeting purine metabolism in cancer cells.

While disrupting purine supply effectively suppresses tumor cell proliferation in experimental models, complete purine depletion in tumor cells is not practical in the clinical setting. Instead, it is more likely that therapies will induce purine shortage rather than total depletion. Thus, elucidating how cancer cells respond to purine shortage, including their adaptive mechanisms and vulnerabilities under such stress, is critical for improving therapeutic designs.In this study we demonstrate that upon purine shortage, cancer cells employ microtubules to assemble complexes resembling purinosomes, in which de novo purine biosynthesis enzymes, along with heat shock proteins (HSP70 and HSP90), form multi-subunit macromolecular complexes as an adaptive mechanism to enhance purine biosynthesis ^35–40^. Additionally, we show that cancer cells utilize 5-Methylthioadenosine phosphorylase (MTAP)-dependent purine salvage pathway as an alternative source of purine supply both in purine-rich conditions, and more prominently, during purine shortage stress when de novo purine biosynthesis is compromised. Consequently, the therapeutic efficacy of DRP-104 can be enhanced by inhibiting MTAP, as the combination of DRP-104 with MTAP inhibition results in superior anti-tumor effects in vivo. Finally, despite the versatility and resilience of cancer cells’ purine supply machinery, we find that purine shortage-stressed tumor cells display higher levels of DNA damage and activation of the cytosolic double-stranded DNA-sensing cGAS-STING-pathway, indicating compromised immune-evasive ability and providing a mechanistic explanation for the previously observed DRP-104-augmented anti-tumor immune response in several tumor models ^10–13^. Collectively, these results highlight cancer cells’ resilience to purine shortage stress while identifying microtubules, MTAP-mediated purine salvage, and impaired immunoevasion as therapeutically targetable vulnerabilities. As microtubules are targets of several commonly used chemotherapeutic drugs and homozygous *MTAP* loss occurs commonly in cancers, our observations may represent a general metabolic mechanism that can be exploited for the treatment of multiple cancer types.

## Materials and Methods

### Sex as a biological variable

Our study exclusively examined male mice since the prostate cancer model is only relevant in males.

### Cell lines and cell cultures

Prostate cancer cell lines (C4-2, PC3, NCI-H660, and TrampC2) and lung cancer cell line NCI-H358 were purchased from the American Type Culture Collection (ATCC) via Duke Cell Culture Facility (CCF). The cell lines’ STR profiles were confirmed by Duke CCF, and the cell lines were tested periodically to ensure that they were mycoplasma-free. All human prostate cancer cell lines were cultured as previously described ^22^. Briefly, C4-2 and PC3 cells were maintained in RPMI 1640 medium with L-glutamine (Gibco, 11875093, ThermoFisher, Waltham, MA, USA), supplemented with 10% fetal bovine serum and 1% penicillin-streptomycin. NCI-H660 cells were maintained in stem cell media (Advanced DMEM/F-12 supplemented with X1 B-27 supplement, 10 ng/mL FGF-2, 10 ng/mL EGF, X1 Glutamax, X1 streptomycin and penicillin). TrampC2 cells were maintained in DMEM with L-glutamine (Gibco, 11965092, ThermoFisher), supplemented with 10% fetal bovine serum, 0.005 mg/ml human insulin, and 10 nM dehydroisoandrosterone and 1% penicillin-streptomycin. NCI-H358 cells were maintained in RPMI 1640 medium (Gibco, 11875093, ThermoFisher), supplemented with 10% fetal bovine serum and 1% penicillin-streptomycin. Glioma CT-2A and GL261 (Redflu-tagged) cell lines (generous gifts from Drs. Darrell Bigner and Vidyalakshmi Chandramohan, Duke University) were cultured as in the previous study ^41^. The *Mtap* knockout CT-2A line was previously described ^42^. The *Mtap*-null derivative GL261 lines were generated following previously described procedure for *Mtap* ^42^. DRP-104 adaptive cells were generated by treating parental cell lines with DRP-104 (generally in the range of 2-4 µM) to obtain populations that ultimately developed adaptation as defined by their continuous propagation in the presence of this prodrug. In general, it takes 3-4 weeks to obtain such “adaptive” cells, which were subsequently maintained in DRP-104-containing media.

### Chemicals and antibodies

DRP-104 and DON were previously described^22^. MTDIA was obtained from MedKoo Biosciences, Inc (cat# 407244) or from MedChemExpress (cat# HY-101496). Antibodies used for immunohistochemistry included the following: CD3 (1:100 dilution; PB9093, BOSTER, USA), CD8⍰ (1:100 dilution; A02236-1, BOSTER, USA), PD-1 (1:100 dilution; A00178, BOSTER, USA), PD-L1 (1:500 dilution; 17952-1-AP, Proteintech, USA), p-IRF3 (Ser396) (1:50 dilution; MA5-14947, Themo Fisher), and γ-H2AX (1:50 dilution; 9718, Cell Signaling Technology). Antibodies used for immunoblotting included the following: PSA (1:1000 dilution; cat# 5365S, Cell Signaling Technology), AR (1:1000 dilution, ab74272, Abcam), MTAP (1:1000 dilution, A20907, ABclonal), p-IRF3 (Ser396) (1:1000 dilution; MA5-14947, Themo Fisher), and γ-H2AX (1:1000 dilution; 9718, Cell Signaling Technology). Detailed information for all reagents used in this study were also provided in **Supplementary Materials**.

### Analysis of public datasets

The Cancer Genome Atlas (TCGA) Prostate Adenocarcinoma (TCGA-PRAD) and SU2C/PCF Dream Team datasets were retrieved from the cBioPortal (http://www.cbioportal.org/) for Cancer Genomics. According to the above datasets, comparative expression and survival analysis were performed with cBioPortal, Gene Expression Profiling Interactive Analysis (GEPIA) and Prostate Cancer Transcriptome Atlas (PCTA) bioinformatic tools.

### In vitro cell growth assays

In vitro cell growth assays were performed following previously described procedure^22,43^. Briefly, adherent cells were plated onto a 96-well plate at ∼5% confluency with at least 3 biological replicates. Cell growth was monitored and quantified by confluency using the IncuCyte S3 Live-Cell Analysis system (Sartorius, Göttingen, Germany) with nine images per well taken with a 20x or 4x objective. The change in confluency was normalized to the confluency at Day 0. All cell growth assays were repeated in at least two independent experiments.

### Metabolite profiling

C4-2/MDVR metabolite profiling data was retrieved from our previous study ^22^. Metabolite profiling of C4-2 and PC3 was performed following previously described procedures ^22^. Briefly, cells were plated into six-well plates at 35% confluency with 3 technical replicates and treated with DMSO (vehicle), 5 μM DRP-104, 6 μM MTDIA, and 5 μM DRP-104 plus 6 μM MTDIA for 48 hours. The confluency of each of the wells was measured by the IncuCyte S3 Live-Cell Analysis system (Sartorius) before metabolite isolation. Cellular metabolites were extracted, as in our previous study ^22,44^. Cells were collected in 80% methanol to extract the metabolites. After centrifugation, the supernatant was dried with a speed vacuum, and dry pellets were reconstituted into 30μL of sample solvent (water: methanol: acetonitrile, 2:1:1), followed by centrifugation at 20,000g for 3 minutes. The supernatant was transferred to LC vials, and 3 μL was submitted to be analyzed with high-performance liquid chromatography-high-resolution mass spectrometry (LC-MS) using previously described procedures ^22,44^.

### Immunofluorescent staining and imaging for purinosome detection

pFGAMS-EGFP plasmid was a gift from Dr. Stephen Benkovic (obtained from Addgene, plasmid^#^ 99107) ^40^. For purinosome detection, cells (2*10^4^ per well) were plated on μ-Slide 4 well chambered coverslip (Ibidi, # 80426, Gräfelfing, Germany) and allowed to attached one day before the transfection. Then, cells were transiently transfected with plasmid pFGAMS-EGFP using Lipofectamine 3000 (Invitrogen, L3000015, ThermoFisher) according to the manufacturer’s instructions. When pertinent, at the time of transfection cells were also transiently treated with agents such as DRP-104, Docetaxel or MTDIA. Two days later after the transfection, cells were washed with PBS, fixed in 4% formaldehyde (Santa Cruz Biotechnology, sc-281692, Dallas, TX, USA) at room temperature for 15 minutes, and permeabilized with 0.2% Triton X-100 (Bio-Rad, # 1610407, Hercules, CA, USA) for 15 minutes. They were then washed and stained with 1 µg/mL DAPI (MilliporeSigma, # D9542, Darmstadt, Germany) before the slides were mounted with ProLong Gold Antifade Mountant (ThermoFisher, # P36394). The slides were imaged with Zeiss 880 Airyscan Fast Inverted Confocal Microscope and analyzed using the Zeiss Zen 3.9 (blue edition) software (Carl Zeiss Microscopy GmbH, Oberkochen, Germany).

For endogenous PAICS immunofluorescent staining, cells were plated and treated as the above procedures without transfection. After 3 days of treatment, cells were washed with PBS, fixed in 4% formaldehyde (Santa Cruz Biotechnology) at room temperature for 15 minutes, and permeabilized with 0.2% Triton X-100 (Bio-Rad) for 15 minutes. Slides with fixed and permeabilized cells were blocked with blocking buffer (1% BSA, 0.2% Triton X-100, 10% goat serum, 0.2 M glycine in PBS) at room temperature for 1 hour, and then incubated with primary antibody (anti-PAICS, 12967-1-AP, Proteintech, Rosemont, IL, USA) diluted in antibody dilution buffer (1% BSA, 0.2% Triton X-100 in PBS; 1:200 dilution) at 4 °C overnight. The next day, slides were washed three times with PBS and incubated with the secondary antibody (anti-rabbit Alexa 594, A-11037, ThermoFisher) 1:500 diluted in antibody dilution buffer at room temperature for 1 hour. Then, cells were washed and stained with 1 µg/mL DAPI (MilliporeSigma) before the slides were mounted with ProLong Gold Antifade Mountant (ThermoFisher). The slides were imaged with Zeiss 880 Airyscan Fast Inverted Confocal Microscope and analyzed using the Zeiss Zen 3.9 (blue edition) software (Carl Zeiss Microscopy GmbH).

### CRISPR-mediated gene knockout

The *MTAP* knockout plasmid was constructed using pLentiCRISPR-E backbone (# 78852, Addgene). The sgRNA sequences for targeting the exon of *MTAP* was designed using the CRISPR program (http://crispor.tefor.net) and described previously (*MTAP* sgRNA sequence #1: GGCTCATCTCACCTTCACGG and sequence #2: CGTTTTAGCTCCCGGGCAGAC; Random control sequence #1: CAGCCACCGCACCGGCGTAA and sequence #2: ATGTTGCAGTTCGGCTCGAT)^45^. In brief, the paired sgRNA were first phosphorylated and annealed using T4 polynucleotide kinase (# M0201S, NEB), followed by ligation with the purified, linearized LentiCRISPRv2E vector (pre-digested with the restriction enzyme BsmBI) using T4 DNA ligase (# M0202S, NEB). Recombinant plasmids were verified to contain expected sgRNA sequences via colony PCR, using LKO.1 5’ and pLentiCRISPR-R1/antisense primers). The *MTAP* CRISPR/Cas9 knockout lentivirus was produced by transfecting HEK293T cells with TransIT-X2® Dynamic Delivery System (# MIR 6004, Mirus Bio) and plasmids including pMD2.G, psPAX2, and two recombinant plasmids at a ratio of 3:4:6. Control lentivirus was prepared in parallel using LentiCRISPRv2E plasmids ligated with the random sequences. *MTAP* and control knockout in vitro model was generated by transducing C4-2 cell lines with lentivirus at a multiplicity of infection (MOI) of 1–2 in a laminin-coated 6-well plate. After 24 hours, the virus-containing medium was replaced with fresh RPMI1640 (CCF). The transduced cells were then selected with puromycin (# P8833; Millipore Sigma) at 1 µg/mL for several days until all non-transduced cells were eliminated. The surviving cells were subsequently maintained in 0.5 µg/mL puromycin. The successful knockout of *MTAP* was confirmed by anti-MTAP immunoblots.

### Protein extraction and immunoblotting

The cell pellet was washed once with pre-chilled PBS and lysed on ice for 30 minutes using RIPA lysis buffer (# R0278; Sigma-Aldrich) supplemented with phosphatase and protease inhibitors (# 1861281; Thermo Fisher). The lysate was sonicated on ice-water bath five times, then centrifuged at 12,000 rpm for 15 minutes at 4°C. The supernatant was collected and mixed with loading buffer (# 161074; Bio-Rad). The mixture was heated at 95-100°C for 5–7 minutes and then loaded onto an SDS-polyacrylamide gel (SDS-PAGE) for electrophoresis, followed by transfer onto an Immun-Blot PVDF membrane (# 1620177; Bio-Rad). The membrane was blocked in TBST containing 5% non-fat milk for 1 hour and then incubated with the primary antibody at 4°C on a shaker for 12–16 hours. After four washes with TBST, the membrane was incubated with an HRP-conjugated secondary antibody diluted at 1:7,000-1:10,000 at room temperature for 1 hour. Following another four washes with TBST, protein detection was performed using SuperSignal West Femto Maximum Sensitivity Substrate (# 34095; Thermo Fisher) and visualized using the iBright Imager (Thermo Fisher) software. Edited whole gel images are available upon request.

### Reverse transcription-quantitative PCR

Total RNA was extracted from the cell pellet using an RNA extraction kit (Cat# R1055; ZYMO RESEARCH) following the manufacturer’s instructions. The RNA concentration and absorbance were measured using a Nanodrop 2000 (Thermo Fisher). RNA samples with a concentration exceeding 50 ng/µL and an A260/A280 ratio above 1.9 were used for reverse transcription to synthesize cDNA (Cat# 4368814; Thermo Fisher). SYBR Green qPCR Master Mix (Cat# 11184ES08; YEASEN) and corresponding primers (Supplementary Materials) were mixed and added to each reaction well, with each target gene analyzed in duplicate. Real-time quantitative PCR (qPCR) was performed on a QuantStudio 3 instrument (Thermo Fisher). The housekeeping gene Actb/ACTB was used for normalization, and the relative expression of target genes was calculated using the 2^^−ΔΔCT^ method. Primers used for PCR were described in the Supplementary Materials.

### ISRE-GFP reporter gene assays

An ISRE-GFP reporter model was established by transducing target cells with lentivirus (Cat# LVP937-P, GenTarget) at a multiplicity of infection (MOI) of 1–2 in the presence of 4 µg/mL polybrene. After 12–24 hours, the culture medium was replaced with fresh DMEM to remove the residual virus. The infected cells were selected using 1 µg/mL puromycin (Cat# P8833; Millipore Sigma) until all uninfected cells died. The selected cells were maintained in the culture medium containing 0.5 µg/mL puromycin. The purified target cells were seeded at a density of 2%-3% in a 96-well plate, and compounds were added according to the experimental protocol. The cells were placed in an IncuCyte system for real-time monitoring, with scheduled daily scans to record the GFP signal area, mean fluorescence intensity, and cell confluency in each well. The total integrated GFP signal intensity (GCU µM²/image) was calculated as the mean signal intensity × signal area/cell confluency. The relative increase in GFP signal was first calculated compared to day 0 and then normalized to the control group to quantify differences in ISRE pathway activation across treatment groups. Representative images were downloaded from the IncuCyte system.

### In vivo tumor models and treatments

The animal study was approved by the Institutional Animal Care and Use Committee (IACUC) of Duke University (A055-22-03). All mice were purchased from the Jackson Laboratory. For TrampC2-derived tumor modes, 24 immune-competent (wild-type), 6-8 week-old C57BL/6J mice were used. The subcutaneous inoculation was performed by injecting 0.5 million cells in PBS and Matrigel (Corning) mixture (1:1) under the right flank skin using a 27 ½ gauge syringe. This injection was performed under brief isoflurane anesthesia on a shaved ∼1 cm^2^ region of the flank. When the tumors reached approximately 100 mm^3^, mice were randomized into six mice per group and treated three times per week, via subcutaneous (s.c.) and intraperitoneal (i.p.) injection for following treatments: (1) Tween 80: Ethanol: Saline (5:5:90) vehicle (s.c. injection) and corn oil vehicle (i.p. injection); (2) Tween 80: Ethanol: Saline (5:5:90) vehicle (s.c. injection) and 5 mg/kg MTDIA (MedKoo Bioscience, 407244, Durham, NC, USA) (i.p. injection); (3) 2 mg/kg DRP-104 (s.c. injection) and corn oil vehicle (i.p. injection), or (4) 2 mg/kg DRP-104 (s.c. injection) and 5 mg/kg MTDIA (i.p. injection). Body weight and xenograft size were measured 3 times a week with calipers by the same scientist throughout the experiment and calculated using tumor volume = length*(width)^2^ *0.5 formula. Mice were sacrificed at 14 days of treatment because the control tumor size met humane end-stage criteria. Syngeneic tumors were dissected and tumor weight/volume was measured. For PC3-derived xenografts and treatments, 24 NSG mice were injected in the right flank with one million PC3 cells in culture media and Matrigel (Corning, 354234, Glendale, AZ, USA) mixture (1:1). Treatments and tumor monitoring were carried out following procedures that were used for the TrampC2 model.

### Hematoxylin and eosin (H&E) staining and immunohistochemical staining

H&E stains and immunohistochemical stains (IHC) were performed following standard procedures as we previously described ^21,43^. Formalin-fixed paraffin-embedded (FFPE) sections at 4-5 mm were deparaffinized and rehydrated. Heat-mediated antigen retrieval was performed in a boiled water bath for 40 minutes in citrate buffer (pH 6.0). Antibodies were incubated at 4 °C overnight. Next day, slides were incubated for 45 minutes at room temperature with HRP-conjugated secondary antibodies (Dako Envision+Kit) and visualized with DAB.

### Quantification and statistical analysis

For quantification of purinosome, cell images were thresholded, and cells with FGAMS or PAICS granules were counted. For each biological repeat, at least 21 cells (for the quantification of FGAMS-EGFP granules, only transfected cells were counted) were analyzed in each group and the percentage of puncta positivity was quantified. Statistical analyses were performed by GraphPad Prism 9. The unpaired two-tailed t-test was used for comparison between two groups, whereas one-way or two-way ANOVA was used for multigroup comparisons.

## Results

### Cancer cells under purine shortage stress reprogram their purine biosynthesis machinery as an adaptive response

Purines provide essential building blocks and serve as energy sources that cancer cells rely on for proliferation. Consequently, purine biosynthesis, particularly the de novo pathway, has been explored as a therapeutic target for treating various cancers, such as pancreatic and brain cancers ^33,45–47^. Purine shortage can be induced by certain anti-cancer drugs, such as Methotrexate, an inhibitor of dihydrofolate reductase (DHFR), although toxicity remains a major limitation ^34^. The recently developed, well-tolerated, clinical-grade prodrug of the broad-acting glutamine antagonist DON, known as DRP-104 (Sirpiglenastat), has shown excellent efficacy in several preclinical cancer models including PCa, and its direct suppressive effects on tumor cells are accompanied by the disruption of purine biosynthesis ^8–10,22^. These findings suggest the feasibility of exploring this well-tolerated prodrug as an effective purine supply-blocking agent for cancer treatment.

Several additional lines of evidence further support this feasibility. First, DON (i.e., DRP-104) inhibits at least 10 enzymes in seven glutamine-utilizing metabolic pathways, with a wide range of potencies (Ki values from ∼µMs to ∼mMs) ^23–32^. Notably, it inhibits FGAMS, an enzyme in the purine synthesis pathway, with the lowest Ki (∼1.1 µM) ^23–32^, suggesting that purine biosynthesis is likely among the pathways most susceptible to DRP-104 (**Supplementary Fig. 1A-B**). Additionally, unbiased metabolic profiling in PCa cell lines PC3 and C4-2 demonstrated that purines were among the most affected metabolites following DRP-104 treatment (**Fig. 1A, Supplementary Fig. 2**). Importantly, supplementation with exogenous purines rescued cancer cells from the suppressive effects of DRP-104 (or DON); and this rescue effect was not unique to PCa cells, as it was also observed in NCI-H358, a lung cancer cell line (**Fig. 1B, Supplementary Fig. 3A**). Gene Set Enrichment Analysis (GSEA) of publicly available gene expression data ^48^ confirmed that the purine biosynthesis pathway is significantly upregulated during PCa pathogenesis **(Supplementary Fig. 3B-C).** Finally, higher expression of FGAMS-encoding PFAS gene or the set of six genes encoding core enzymes of the purine biosynthesis machinery, namely purinosomes^36^, was correlated with worse survival in PCa patients **(Supplementary Fig. 3D-F)**, providing further rationale for targeting purine supply as a therapeutic strategy.

**Fig. 1.**
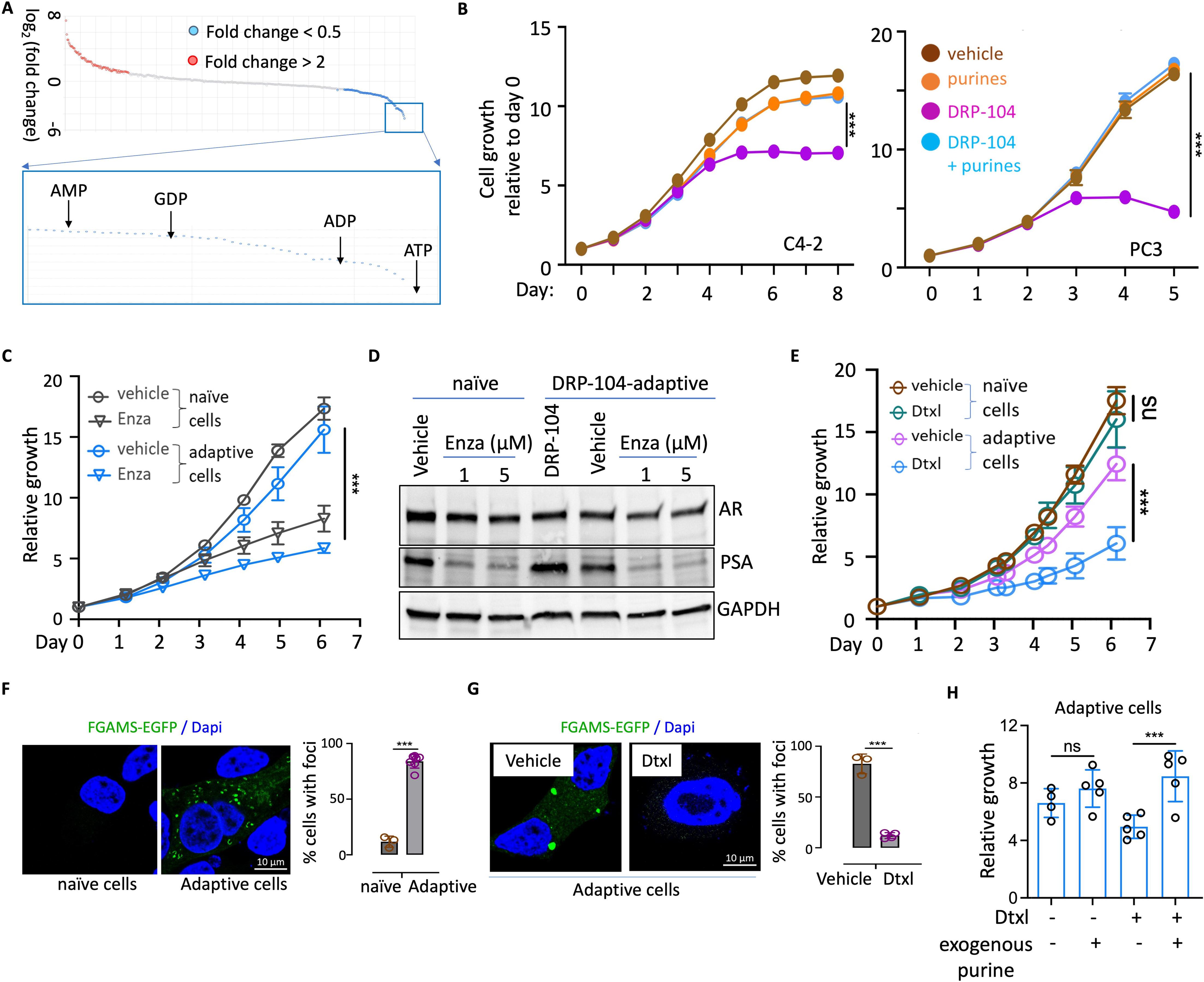
Prostate cancer cells stressed by purine shortage reprogram their purine biosynthesis machinery as an adaptive response. **(A)** PC3 cells were treated with DRP-104 and subjected to metabolic profiling. Analysis of the altered metabolites in DRP-104-treated cells vs. the control-treated cells showed that purines were among those that were most strongly affected. **(B)** Tumor cells were treated with indicated agents and the relative cell proliferation was monitored by IncuCyte. For DRP-104, 1 µM (for C4-2) or 2 µM (for PC3) were used. Exogenous purines used included adenosine plus guanosine (each at concentration ranging from 30-100 µM). Data are depicted as mean ± SD. **(C)** Treatment-naïve (parental) or DRP-104-adaptive C4-2 cells were treated with Enzalutamide (Enza) and cell proliferation was determined. Similar results were obtained in LNCaP cell lines treated with hormone depletion (by using charcoal-stripped FBS). Data are depicted as mean ± SEM. **(D)** Cells used in (C) were subjected to immunoblot assays. **(E)** Treatment-naïve or DRP-104-adaptive C4-2 cells were treated with vehicle or with low doses of Docetaxel (Dtxl) (e.g., 0.1 nM) and the relative cell proliferation was determined. ***p<0.001. **(F)** Treatment-naïve or DRP-104-adaptive C4-2 cells were subjected to purinosome formation assays using FGAMS-EGFP as the biomarker. (**G**) DRP-104-adaptive C4-2 cells were treated with vehicle or with Docetaxel (Dtxl, 1 nM) transiently and the abundance of purinosomes was determined. Representative images and the quantification were shown. Similar results were obtained in PC3 cell lines. **(H)** DRP-104-adaptive C4-2 cells were treated with indicated agents and their relative growth was shown (day 4 data). ***p<0.001.

While DRP-104 effectively suppresses tumor cell proliferation, it is unlikely to completely deplete purines from tumor cells. Instead, it is more feasible to induce a state of purine shortage. Over time, cells that survive this purine shortage may adapt to the stress and ultimately resume propagation, as observed with many targeted therapies. This hypothesis is supported by the observation that while DRP-104 or DON (typically in the range of 2-4 µM) effectively suppressed tumor cell proliferation, the treatment did not eradicate tumor cells. Eventually, the cells adapted to this stress and resumed growth, leading to the emergence of DRP-104-adaptive (resistant) cell lines capable of proliferating in the presence of DRP-104 (**Fig. 1C**, **Supplementary Fig. 4**). Although DRP-104-adaptive PC3 cells exhibited substantially slower growth compared to their DRP-104-naïve (parental) counterparts, DRP-104-adaptive C4-2 cells propagated at a rate similar to that of their treatment-naïve counterparts (**Fig. 1C**, **Supplementary Fig. 4**). We thus utilized the C4-2 models to investigate the mechanisms underlying the adaptation. Since C4-2 cells are AR-dependent and respond to AR inhibition, we first evaluated the response of these purine shortage-adaptive derivatives to the AR antagonist enzalutamide. These cells retained their susceptibility to the AR signaling blockade (**Fig. 1C-D**, **Supplementary Fig. 4**), consistent with the expectation that purine dependency is not exclusive to AR-driven cell proliferation. This also suggests that prior DRP-104 exposure does not interfere with the effectiveness of subsequent AR-targeting therapies. Most notably, we found that, compared to their DRP-104 naïve counterparts, DRP-104-adaptive cells displayed increased sensitivity to Docetaxel (**Fig. 1E**), a microtubule-stabilizing chemotherapeutic drug commonly used for advanced and metastatic PCa ^49–54^.

Microtubules, as key components of the cellular cytoskeleton, play essential roles in various cellular processes. In the context of cancer treatments, the focus has traditionally been on their roles in cell division, while other cellular processes have received less attention. One such underexplored process is purine biosynthesis in mammalian cells under purine shortage stress. In response to this stress, cells assemble multiple-subunit macromolecular complexes of de novo purine biosynthesis enzymes, known as purinosomes, as an adaptive mechanism to more efficiently ramp up purine biosynthesis^35–40^. Microtubules are essential for the assembly and proper spatial distribution of purinosomes, as demonstrated in previous studies^36,39,55–57^. Several lines of evidence suggest that the heightened sensitivity of DRP-104-resistant cells to Docetaxel, a microtubule-targeting drug, is due to their reprogrammed purine biosynthesis machinery and acquired dependency on purinosomes. First, using EGFP-tagged FGAMS, a well-established marker for purinosome detection ^35–40^, we observed a significant increase in purinosome-resembling foci in the resistant/adaptive cells, implicating purinosomes in enabling PCa cell resistance and adaptation (**Fig. 1F**). Secondly, treatment with Docetaxel - at doses that did not affect overall cell proliferation or survival – led to a reduction in purinosome abundance in the resistant cells, consistent with the drug’s microtubule-targeting effects (**Fig. 1G**). Additionally, exogenous purine supplementation protected the resistant cells from the suppressive effects of Docetaxel (**Fig. H**), further underscoring the link between purinosome dependency and drug sensitivity.

Further supporting the role of DRP-104 as an inducer of purine shortage - and consistent with the findings from the DRP-104-adaptive cells described above - subsequent experiments confirmed that transient DRP-104 treatment effectively induces the formation of purinosomes across multiple cancer cell lines, including PCa cell lines (e.g., PC3 and C4-2), a lung cancer cell line (NCI-H358), and a mouse glioma cell line CT-2A (**Fig. 2A-B**). Results from the exogenously expressed EGFP-FGAMS-based purinosome detection were further validated by immunofluorescent staining targeting endogenous PAICS, another key enzyme in the de novo purine biosynthesis pathway and a commonly used marker for purinosome assays ^35–40^ (**Fig. 2C-D**).

**Fig. 2.**
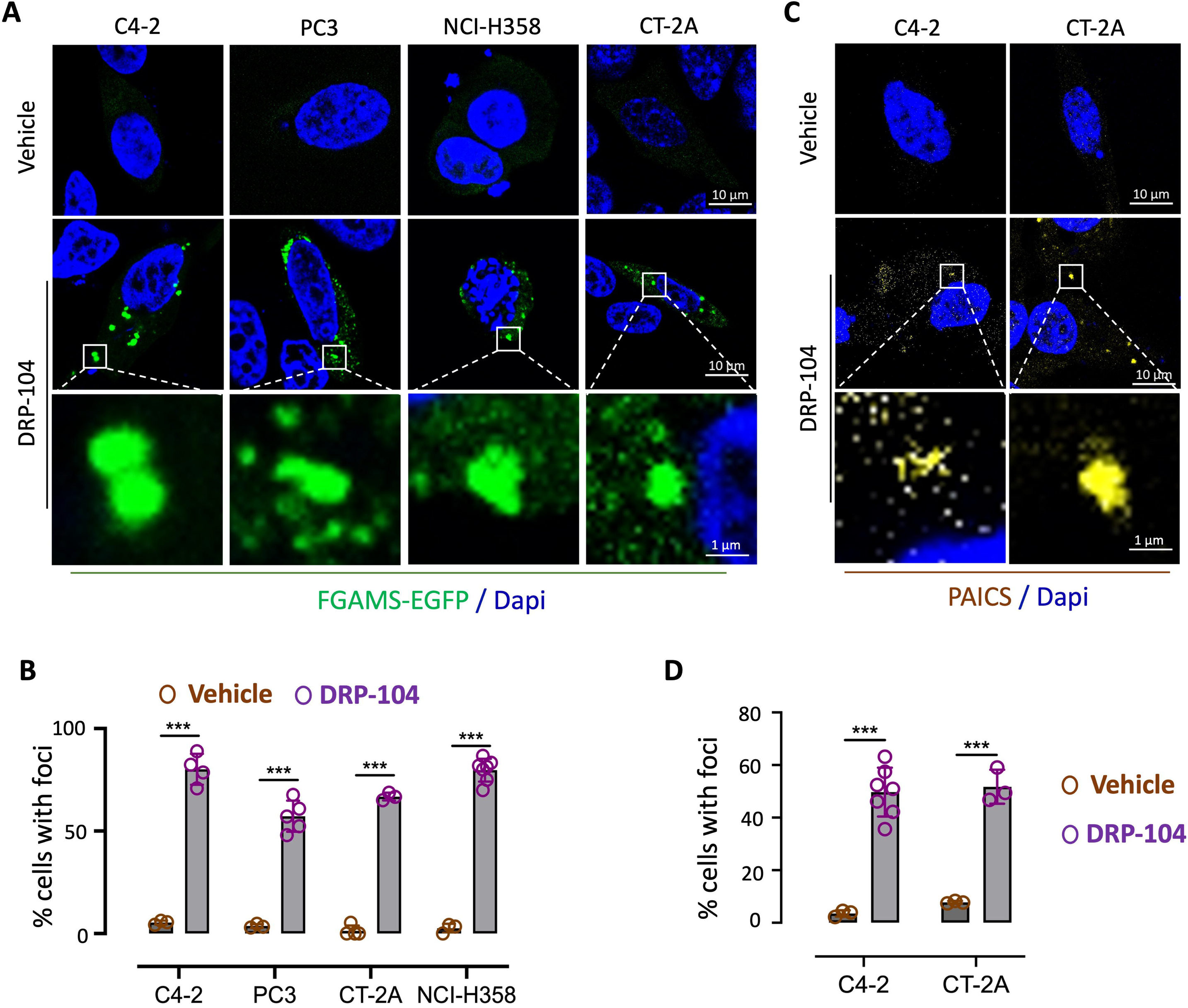
Cancer cells respond to DRP-104 treatment by promptly forming purinosomes. Cancer cell lines were transfected with exogenous FGAMS-EGFP expressing plasmid and subjected to DRP-104 treatment (day 0), and the presence of purinosomes (FGAMS-EGFP foci) were determined (day 2). Alternatively, tumor cells were treated with DRP-104 (day 0), and the presence of purinosomes was detected by anti-PAICS immunofluorescent staining and confocal microscopy analysis (day 2). Representative images (**A, C**) and quantifications (**B, D**) were shown. ***p<0.001.

Collectively, these results suggest that cancer cells under purine shortage stress reprogram their purine biosynthesis machinery – particularly through the microtubule-mediated assembly of purinosomes – as an adaptive mechanism.

### MTAP-dependent purine salvage serves as a critical source of purine supply in cancer cells

Purine biosynthesis comprises two primary pathways: de novo biosynthesis and the salvage process. In the purine salvage pathway, MTAP converts 5′-methylthioadenosine (MTA), a byproduct of the polyamine biosynthesis pathway, into methionine and adenine. The adenine is subsequently converted by Adenine Phosphoribosyltransferase (APRT) to form AMP **(Supplementary Fig. 1B)**. Understanding the role of MTAP-mediated purine salvage in PCa cells’ response to DRP-104 is critical. While MTAP alterations are prevalent in some cancer types, such as glioblastoma where approximately half of all tumors harbor homozygous MTAP loss, they present as various types of alterations in PCa and occur in only ∼2-3% of cases (**Supplementary Fig. 5A**). Previous studies have investigated MTAP inhibition as a strategy to suppress multiple cancer types ^58–60^. Inhibition of MTAP has been shown to directly suppress PCa cells ^61^, and that its inhibition along with stimulation of polyamine biosynthesis exacerbates the stress of one-carbon unit loss, thereby suppressing PCa progression^62^. However, the extent to which MTAP-dependent purine salvage contributes to purine supply in PCa remains unclear, especially in the context of cellular adaptation to purine shortage. To address this, PCa cells were treated with MTDIA, a specific transition state analogue inhibitor of MTAP ^58–60^, followed by unbiased metabolic profiling analysis. The results demonstrated that MTAP inhibition predominantly affected purine metabolism (**Fig. 3A**). Further, treating prostate cancer cell lines (C4-2, PC3) and the lung cancer cell line NCI-H358 with MTDIA resulted in an increased presence of purinosomes (**Fig. 3B-C, Supplementary Fig. 5B**), indicating that MTAP inhibition induces purine shortage stress. Additional experiments using CT-2A cell line models, including an *Mtap*-null derivative line^42^, revealed a higher abundance of purinosomes in parental (*Mtap*-intact) CT-2A cells treated with MTDIA and in *Mtap*-null CT-2A cells compared to their parental, vehicle-treated counterparts (**Fig. 3D-E, Supplementary Fig. 5C**).

**Fig. 3.**
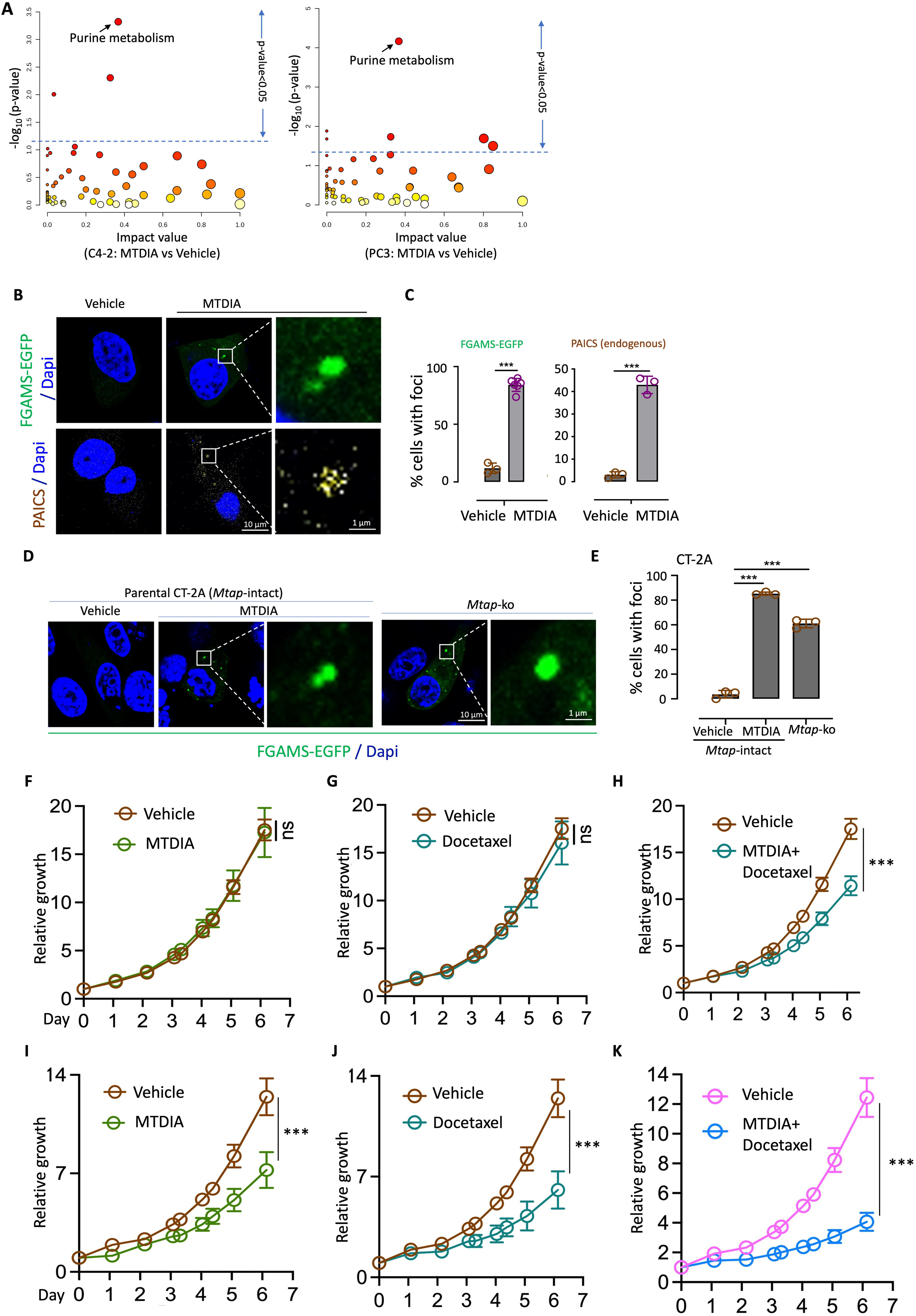
MTAP-dependent Purine salvage is a critical source of purine supply in tumor cells. **(A)** Prostate cancer cells were treated with DRP-104 (4 µM, 2 days) and subjected to metabolic profiling to identify metabolites that were affected. Pathway topology analysis was performed to identify metabolism pathway that were significantly altered. **(B-C)** C4-2 cells expressing exogenous FGAMS-EGFP were subjected to MTDIA treatment (day 0), and the presence of purinosomes was detected (day 2). Alternatively, cells were treated with MTDIA (day 0), and the presence of purinosomes was detected by anti-PAICS immunofluorescent staining and confocal microscopy analysis (day 2). Representative images and quantifications were shown. **(D-E)** CT-2A cells transfected with exogenous FGAMS-EGFP expressing plasmid and subjected to MTDIA treatment (5 µM, day 0), and the presence of purinosomes was detected (day 2). Separately, *Mtap*-ko CT-2A cells were used for similar FGAMS-EGFP-based purinosome detection. Representative images (D) and quantifications (E) were shown. **(F-H)** Treatment-naïve (parental) and **(I-K)** DRP-104-adaptive C4-2 lines were treated with vehicle, MTDIA (3 µM), Docetaxel (0.1 nM) or MTDIA plus Docetaxel and cell growth was determined. Data are depicted as mean ± SEM. ***p<0.001.

While it is generally believed that tumor cells primarily rely on *de novo* purine biosynthesis while also recycling purine precursors from the microenvironment ^63^, the results presented here suggest that cancer cells, even when cultured under standard conditions, sense purine shortage upon loss of MTAP function. Intriguingly, despite the clear role of MTAP-dependent purine salvage as a source of purine supply, cancer cell proliferation was not affected by MTAP inhibition, as shown in previous findings in glioma models ^45^, as well as in prostate cancer models described here (**Supplementary Fig. 5D**). In contrast, cancer cells adaptive to DRP-104 displayed measurable susceptibility to MTAP inhibition (**Supplementary Fig. 5D**).

The increased abundance of purinosomes, which are capable of more efficient de novo purine biosynthesis^35^, in the MTAP-deficient cells led us to hypothesize that forming purinosomes is a way for tumor cells to compensate for the loss of purine salvage. In agreement with this hypothesis, we found that treating cancer cells with MTDIA in the presence of Docetaxel, at a dose that did not affect cell proliferation (**Fig. 3G**), resulted in significant suppression of proliferation (**Fig. 3H**). Most remarkably, similar treatment experiments in DRP-104-adaptive cells, which were more susceptible to Docetaxel compared to their naïve counterparts (as shown in **Fig. 1E**, and again comparing **Fig. 3J** versus **3G**), showed that compared to their naïve counterparts, these cells were also more sensitive to MTDIA (**Fig. 3** versus **3FI**), and displayed even higher susceptibility to the combinational treatment of MTDIA plus Docetaxel (**Fig. 3K** versus **3H**).

Collectively, these results suggest that MTAP inhibition triggers adaptive stress responses in cancer cells, marked by purinosome formation, thus underscoring the critical role of MTAP-dependent purine salvage as a consequential source of purine supply for cancer cells. Additionally, these findings corroborate the previously proposed critical roles of microtubule-mediated formation of purinosomes and suggest a functional link between MTAP and microtubules in the context of purine supply and shortage. Upon the loss of MTAP-mediated purine salvage, tumor cells resort to microtubules to assemble purinosomes (for more efficient purine biosynthesis as a compensatory adaptation), and vice versa. These two compensatory mechanisms of purine supply together become essential for enabling tumor cells to adapt to compromised de novo purine biosynthesis, as in the case of DRP-104 treatment.

### Inhibiting MTAP sensitizes tumor cells to DRP-104 treatments

Several studies have investigated MTAP inhibition as a cancer treatment strategy using MTDIA, an orally bioavailable, potent, and specific transition state analogue inhibitor of MTAP ^58–62^. The feasibility of therapeutically targeting MTAP, along with the compensatory relationship between MTAP-dependent purine salvage and de novo purine biosynthesis described above, led us to test the hypothesis that (1) MTAP-mediated purine salvage protects cancer cells from DRP-104-induced purine shortage and (2) inhibiting MTAP can enhance the tumor-suppressive effect of this prodrug.

First, we treated PCa cell lines with DRP-104 in the presence of exogenous MTAP substrate, 5’-deoxyadenosine (5’dAdo), and observed that 5’dAdo mitigated DRP-104-induced suppression of cell growth (**Fig. 4A, Supplementary Fig. 6A**). This finding suggests that although the strength of the purine salvage pathway may vary across cancer cell lines, excess MTAP substrate is protective. In contrast, the presence of MTAP inhibitor MTDIA potentiated the suppressive effects of DRP-104 (**Fig. 4B, Supplementary Fig. 6B**). We confirmed that the protective and damaging effects of 5’dAdo and MTDIA, respectively, were not due to their influence on the conversion of the prodrug, as similar effects were observed when using DON (**Fig. 4C**). To further corroborate these findings, we compared the metabolic profiles in C4-2 cell lines treated with DRP-104 plus MTDIA to those of the cells that were treated with DRP-104 alone. This analysis revealed that the combination treatment primarily affected purine metabolism, along with cysteine and methionine metabolism pathways, consistent with MTDIA-induced disruption of the purine supply (**Supplementary Fig. 6C-D, Supplementary Table 1**). As expected, purinosomes were prominently present in tumor cells that were treated with DRP-104 plus MTDIA, confirming both the occurrence of purine shortage in these cells and the enhanced suppressive effect of the DRP-104 and MTDIA combination (**Supplementary Fig. 6E**).

**Fig. 4.**
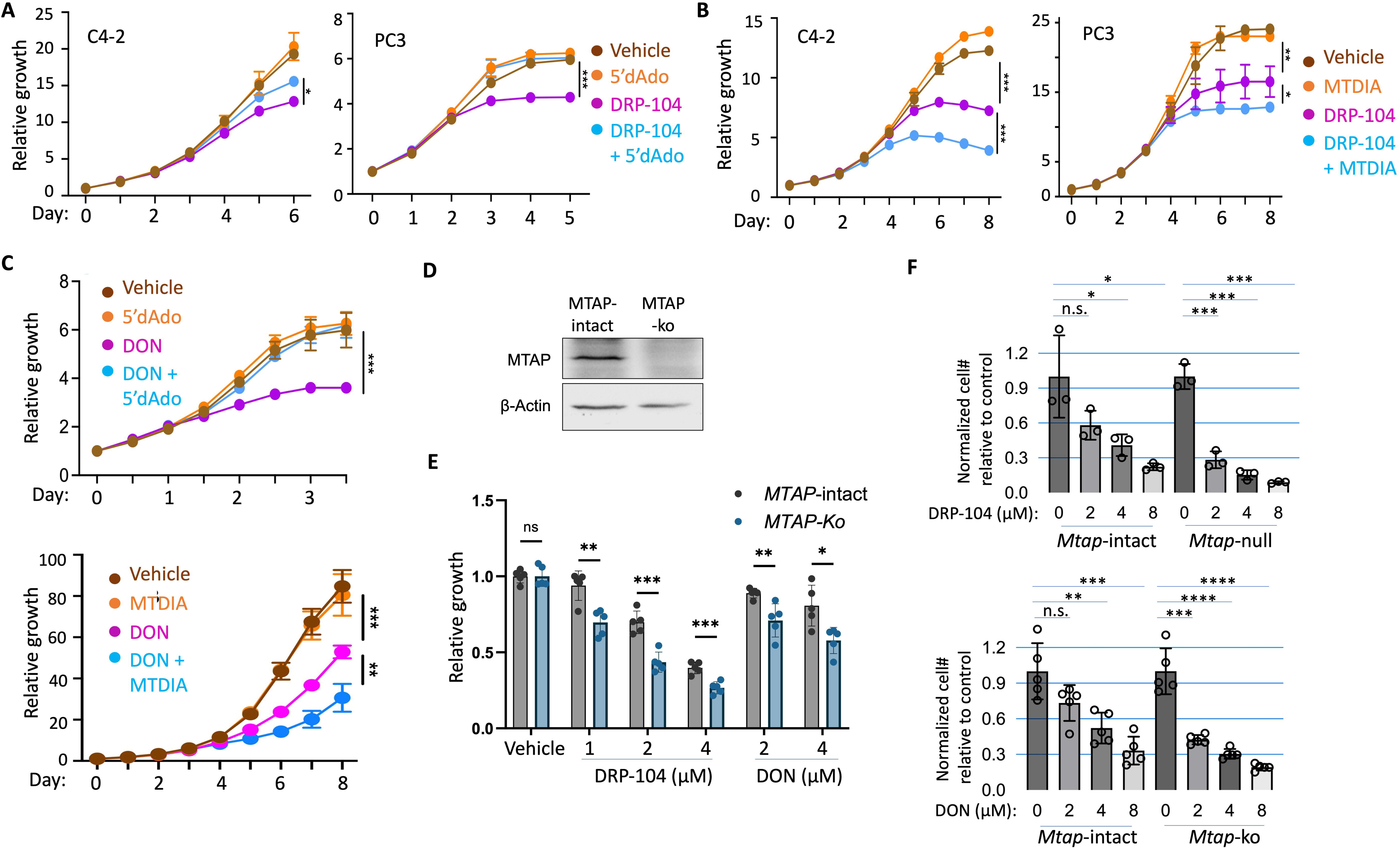
MTAP deficiency sensitizes tumor cells to DRP-104 treatments. **(A-B)** Tumor cells were treated with indicated agents and the relative cell proliferation was monitored by IncuCyte. 2 µM of DRP-104 was used for all cell lines; concentrations of 5’dAdo used varied from 10-60 µM, yielding consistent results; concentration of MTDIA used was 3 µM. Data are depicted as mean ± SD. **(C)** PC3 cells were treated with indicated agents and their growth was determined by IncuCyte (4, 30 and 3 µM was used for DON, 5’dAdo and MTDIA, respectively). Data are depicted as mean ± SD. **(D)** Immunoblots confirmed the knockout of *MTAP* in C4-2 cell line. **(E)** C4-2-derived CRISPR control (*MTAP*-intact) and *MTAP*-ko cell lines were treated with vehicle control or with DRP-104 or DON, and relative cell growth (compared to the vehicle control) on day 6 post treatment was shown. **(F)** *Mtap*-intact (parental) and *Mtap*-null CT-2A lines were treated with DRP-104 or DON and their growth was determined (relative cell growth on day 6 and day 5 were shown for DRP-105 and DON treatments, respectively; standard deviations were denoted). For the CT-2A cell lines, exogenous 5’dAdo (10 µM) was supplied in the media to ensure sufficient MTAP substrate was present. *p<0.05, **p<0.01, ***p<0.001.

Furthermore, as a complementary strategy to the chemical approach (i.e., the use of MTDIA), we generated *MTAP*-null derivative cell line from C4-2 using CRISPR-mediated *MTAP* knockout (**Fig. 4D**) and confirmed that the *MTAP*-null cell line was more susceptible to DRP-104 or DON compared to the MTAP-intact controls (**Fig. 4E**). To assess whether MTDIA’s ability to enhance DRP-104 efficacy extends beyond PCa, we tested DRP-104 sensitivity in NCI-H358, a lung cancer cell line. As observed in prostate cancer cell lines, MTDIA alone did not affect cell propagation but increased NCI-H358 cells’ susceptibility to DRP-104 (**Supplementary Fig. 6F**). We further tested a previously described *Mtap*-null cell line derived from mouse glioma cell line CT-2A^42^, and confirmed its increased susceptibility to DRP-104 (or DON) compared to its *Mtap*-intact counterpart (**Fig. 4F**). The role of Mtap in modulating tumor cell susceptibility to purine shortage – induced by treatments such as DRP-104 or DON - was also observed in another glioma cell line, GL261, using both chemical (MTDIA) and genetic (*Mtap* knockout) approaches (**Supplementary Fig. 6G-H**).

Collectively, these findings underscore the protective effects of MTAP-mediated purine salvage in purine shortage induced by DRP-104 therapy. They suggest that *MTAP* status should be a consideration when designing purine blockade-based treatments, and highlight the potential of MTAP inhibitors, such as MTDIA, to enhance the efficacy of purine shortage inducers such as DRP-104 in *MTAP*-intact cancers.

### Combining DRP-104 with MTDIA yields superior therapeutic efficacy in a prostate cancer model in vivo

We next evaluated the therapeutic effects of simultaneous targeting MTAP and de novo purine biosynthesis by combining MTDIA and DRP-104 in vivo. In addition to directly suppressing tumor cells, DRP-104 has been shown to enhance anti-cancer immunity across several cancer types ^10–12^. Notably, previous studies suggested that in certain cancers, such as pancreatic cancer and glioma, *MTAP* loss and the resulting accumulation of its substrate, methylthioadenosine (MTA), contribute to an immunosuppressive tumor microenvironment ^42,64^. These findings raise the possibility that while combining MTDIA with DRP-104 may enhance direct tumor cell suppression, as demonstrated in vitro, any potential immunosuppressive effects of MTDIA might counteract this therapeutic benefit.

To evaluate this possibility, we generated subcutaneous tumors from PCa cell line TrampC2 in immune-competent C57BL/6 host mice to test the effects of the combination treatment versus single-agent treatments. Our previous study showed that the regimen of 2 mg/kg daily DRP-104 treatment, five days per week, effectively suppressed NCI-H660-derived xenograft growth ^22^. To evaluate the therapeutic efficacy of combining DRP-104 and MTDIA, we reduced the treatment frequency as follows: DRP-104 (2 mg/kg), MTDIA (5 mg/kg), or both (2 mg/kg and 5 mg/kg, respectively) were administered three times per week for a total of seven treatments and tumor progression was monitored (see **Supplementary Fig. 7A**). The experiment confirmed that DRP-104 alone had measurable efficacy as expected, while MTDIA alone had minimal effects, as assessed by tumor volume and weight (**Fig. 5A-C**). Remarkably, the combination of both agents resulted in superior tumor suppression (**Fig. 5A-C**), without significantly affecting the body weights of the recipient mice (**Supplementary Fig. 7B**). H&E staining revealed that while tumors from vehicle- and MTDIA-treated mice have sheets of viable tumor cells that were actively proliferating as reflected by frequent mitotic figures (**Fig. 5D**). In contrast, DRP-104-treated tumors had more sparse tumor cells, segregated by immune cells, fibrosis and tumor necrosis (**Fig. 5D**). Notably, residual tissue masses from mice treated with the DRP-104 and MTDIA combination were composed of immune cells and fibrosis without appreciable tumor cells (**Fig. 5D**).

**Fig. 5.**
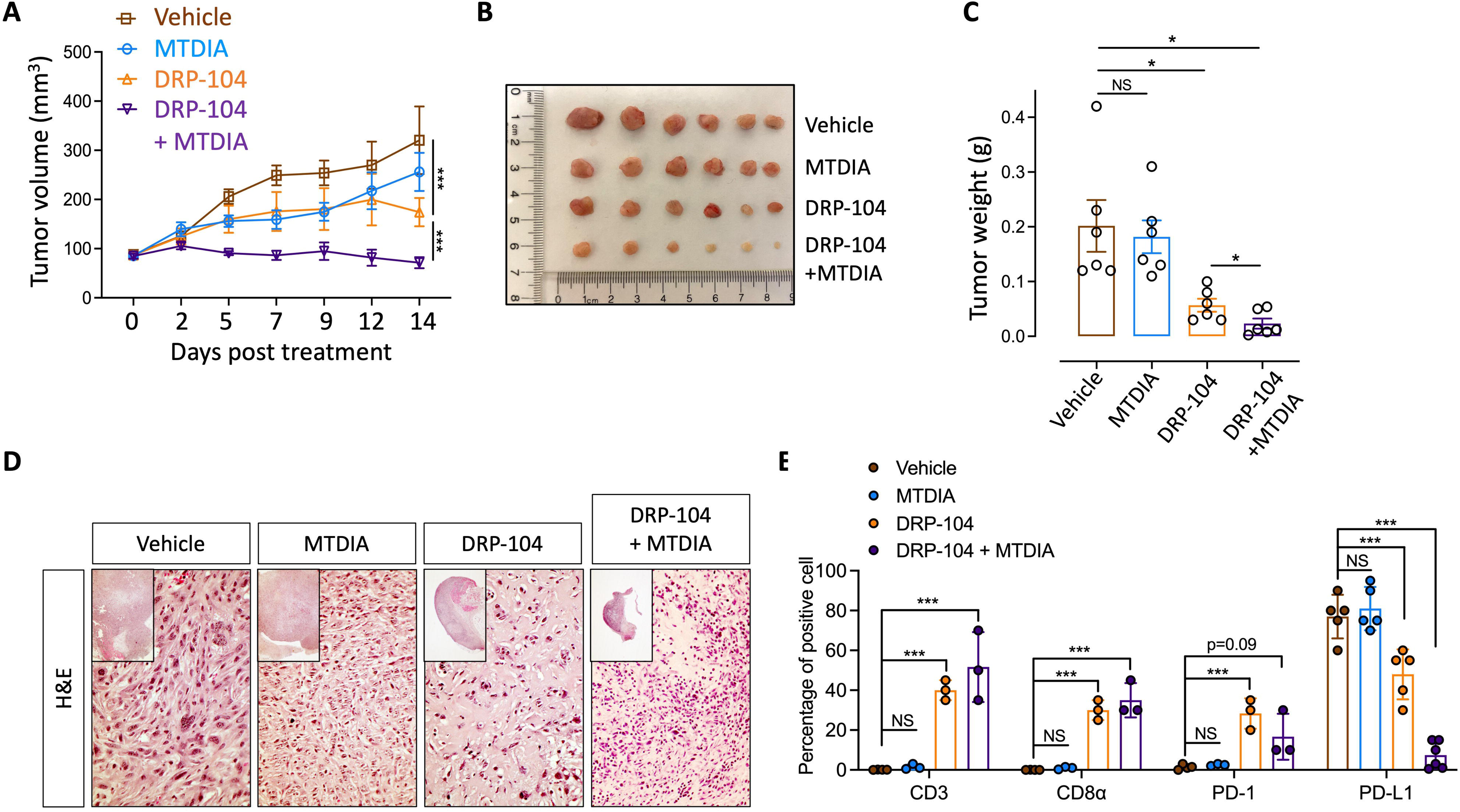
Combining DRP-104 with MTDIA yields superior therapeutic efficacy in an immunocompetent tumor model. **(A-C)** TrampC2-derived tumor-bearing wildtype C57BL/6 male mice were treated with indicated agents and tumor volumes were determined during the two weeks’ treatment period (n=6 mice per arm). Data are depicted as mean ± SEM in (A). Images and weights of tumors at Day 14 (after 6 treatments) were shown (B) and (C). **(D)** Representative images of tumors that were harvested after two weeks’ treatments. **(E)** IHC staining for CD3, CD8α, PD-1 and PD-L1 in the harvested tumors. ns: no significant, *p<0.05, **p<0.01, ***p<0.001.

We then performed immunohistochemical (IHC) assays for selected immune biomarkers, including T lymphocyte markers (CD3 and CD8α) and immune checkpoint markers (PD-1 and PD-L1), to evaluate the effects of the comprehensive purine blockade (i.e., combination of DRP-104 and MTDIA) on the tumor immune microenvironment. These IHC assays revealed that vehicle-treated and MTDIA-treated tumors displayed minimal staining for CD3, CD8α, and PD-1 but high levels of PD-L1, indicating low T cell infiltration and an immunosuppressive tumor microenvironment (**Fig. 5E, Supplementary Fig. 7C**). In contrast, both DRP-104-treated and DRP-104 plus MTDIA-treated tumors showed robust staining for CD3 and CD8α, suggesting strong T cell infiltration (**Fig. 5E, Supplementary Fig. 7C**). Compared with vehicle-treated and MTDIA-treated tumors, DRP-104-treated tumors had significantly reduced PD-L1^+^ cells and an increased presence of PD-1^+^ cells, possibly indicating negative immunoregulatory feedback in tumors whose progression had been mitigated. Most notably, DRP-104 plus MTDIA-treated tumors showed a slight increase in PD-1^+^ cells and were largely devoid of PD-L1^+^ cells (**Fig. 5E, Supplementary Fig. 7C**), consistent with the histological observation that tumor cells were essentially all eradicated in tissues from the combination therapy group (**Fig. 5D**).

The above results suggest that the superior therapeutic efficacy of the combination therapy is likely due to two factors: more effective direct tumor cell-autonomous suppression and the reshaping of the tumor immune microenvironment to stimulate anti-tumor immunity. This implies that the combination therapy may enhance, rather than diminish, DRP-104-induced anti-tumor immunity. Additionally, the findings from the in vivo model further corroborate results from in vitro studies and highlight the potential of combining MTAP inhibitors, such as MTDIA, with DRP-104 to achieve superior therapeutic outcomes in cancers that retain functional MTAP.

### Purine supply deficiency leads to activated cGAS-STING signal pathway

The DRP-104-induced changes in the tumor immune microenvironment prompted us to investigate the mechanisms underlying the effects of this tumor cell-specific prodrug. Several previous findings are particularly noteworthy. First, an abundant supply of purine is critical for tumor cells’ response to DNA damage ^65–67^. Additionally, a compromised DNA damage response is linked to the cytosolic double-stranded DNA-sensing cGAS-STING pathway and innate immunity^68^. Indeed, it has been shown that a deficiency in the DNA damage response in PCa activates the cGAS-STING pathway ^69^. Given that the status of the cGAS-STING pathway is a key determinant of the PCa immune microenvironment ^70^, we assessed the link between DRP-104-induced purine shortage, DNA damage and the cGAS-STING pathway. Intriguingly, transient in vitro treatment with DRP-104 led to elevated levels of _H2AX and phosphorylated IRF3 (p-IRF3), a downstream mediator of the cGAS-STING signal (**Supplementary Fig. 8A**), along with increased expression of interferon response genes such as *Ifnb1* and *Ccl5* (**Supplementary Fig. 8B**). These findings suggest a connection between purine shortage, DNA damage, and activation of the cGAS-STING pathway. As the tissue masses remaining after the treatments were mostly depleted of cancer cells and infiltrated with immune cells in the immune-competent tumor models mentioned above, we sought to establish xenografts in immunocompromised mice to further investigate. Our recent study shows that prostate cancer, together with brain tumors, is among the cancer types where the *STING1* gene is most heavily epigenetically silenced ^71^. Nevertheless, the cGAS-STING pathway remains intact and can be functionally activated in certain prostate cancer cell lines, including PC3 and C4-2^69^. We therefore generated PC3-derived xenografts and treated tumor-bearing immunocompromised mice following the regimens described above (**Supplementary Fig. 7A**). The results corroborated findings from the TrampC2 model, demonstrating the superior efficacy of combining DRP-104 and MTDIA compared to each monotherapy (**Fig 6A**). We harvested the treated tumors at their endpoints and performed H&E staining and IHC for Ki67 to confirm the presence of tumor cells (**Supplementary Fig. 8C**), consistent with the expected direct suppressive effects on tumor cells rather than robust tumor cell eradication through anti-tumor immunity. Most importantly, H2AX and p-IRF3 IHC assays revealed that tumors treated with DRP-104 or combination therapy displayed elevated levels of both markers (**Fig. 6B-C**), supporting a connection between purine shortage, DNA damage, and the cGAS-STING pathway in vivo.

**Fig. 6.**
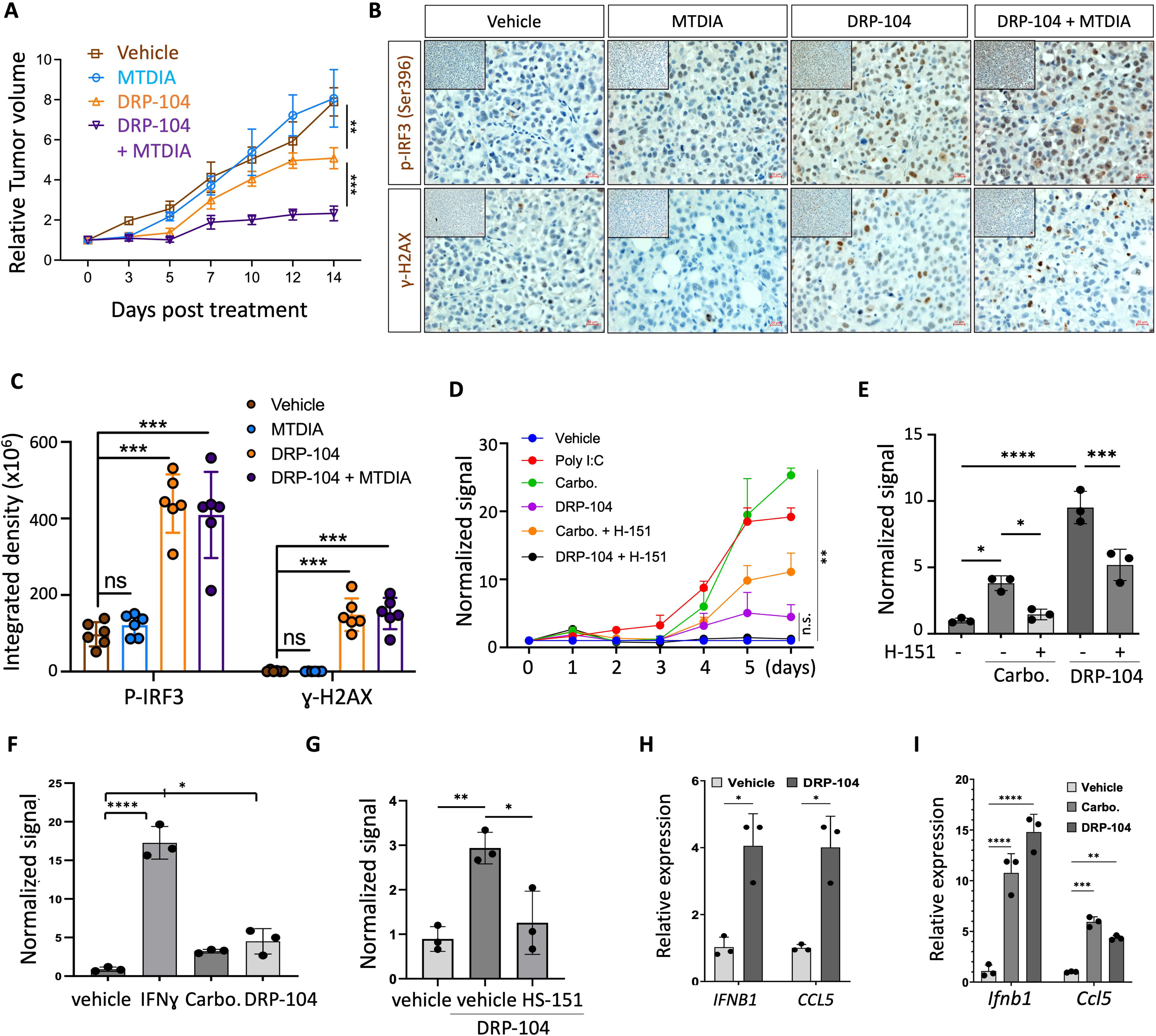
DRP-104 treatments result in activated cGAS-STING pathway. **(A)** PC3-derived tumor bearing mice were treated with indicated agents and tumor volumes were monitored during the treatment period. Data are depicted as mean ± SEM. **(B-C)** PC3-derived xenografts treated with vehicle or with DRP-104 (with or without MTDIA) were used for anti- H2AX and anti-p-IRF3 IHC and (B) representative images and (C) quantifications were shown. **(D)** ISRE reporter-tagged PC3 cells were treated with indicated agents and the reporter signal was monitored by IncuCyte (poly I:C, a known stimulator of the ISRE activity, was included as a positive control). Integrated fluorescent reporter signaling was normalized by cell confluences as determined by IncuCyte. **(E)** ISRE reporter-tagged C4-2 cells were treated with indicated agents and the reporter signal was determined. **(F-G)** ISRE reporter-tagged, *Mtap*-null derivative CT-2A cell line (F) and parental (*Mtap*-intact) CT-2A line (G) were treated with indicated agents and the reporter signal was monitored by IncuCyte (IFN was included as a positive control; day 4 data was shown in F). **(H)** C4-2 cells were treated with indicated agents and the expression of marker genes in the cGAS-STING pathway, *IFNB1* and *CCL5*, was determined. (I) *Mtap*-null CT-2A cells were treated with indicated agents and the expression of marker genes in the cGAS-STING pathway, *Ifnb1* and *Ccl5*, was determined. carbo.: carboplatin. n.s.: not significant, *p<0.05, **p<0.01, ****p<0.0001.

We further employed an ISRE (interferon-stimulated response element) reporter-based assay^72^ to assess the effects of DRP-104 on the cGAS-STING pathway in PC3 cells in vitro. We found that carboplatin, a genotoxic drug that is used for treating CRPC or small cell carcinoma of the prostate, activated the reporter signal as expected, and notably, DRP-104 treatment similarly stimulated the reporter signal, albeit to a lesser degree (**Fig. 6D, Supplementary Fig. 8D**). Furthermore, both carboplatin and DRP-104-stimulated reporter expression were mitigated by H-151, a STING antagonist (**Fig. 6D, Supplementary Fig. 8D**). These findings from the reporter-based assays were corroborated in the C4-2 cell line (**Fig. 6E**). To test whether the DRP-104 and cGAS-STING link is present in other tumor models, we extended the assay to glioma cell line CT-2A, given that the critical roles of purine supply in DNA damage repair have been well established in gliomas^66,67^. We confirmed that purine shortage, induced by either DRP-104 (in the *Mtap*-null CT-2A derivative line) or DRP-104 plus MTDIA (in the parental, *Mtap*-intact CT-2A cells), similarly resulted in activation of the reporter signal (**Fig. 6F-G, Supplementary Fig. 8E-F**). Finally, we used *IFNB1* and *CCL5* as the ISRE-regulated marker genes and assessed their expression in response to the treatments. In PC3 cells, these genes had very low basal expression levels, leading to variable results. However, in C4-2 and in CT-2A models, the expression of these genes was consistently stimulated by DRP-104 treatments (**Fig. 6F-G, Supplementary Fig. 8G**).

Collectively, these results are consistent with the scenario in which purine shortage, such as treatment with DRP-104, leads to activation of the cGAS-STING signal pathway in tumor cells. While the contribution of this pathway’s activation to alterations in the tumor immune microenvironment and adaptive anti-tumor immunity remains to be elucidated, these findings suggest that purine shortage stress compromises the immunoevasive capacity of tumor cells. They also offer a potential mechanistic explanation for the DRP-104-induced changes in the tumor immune microenvironment in the TrampC2 model described above, as well as in tumor models of several other cancer types ^10–13^.

## Discussion

In this study, we demonstrate that purine supply represents a major liability in cancer cells that can be exploited for therapeutic purpose. We show that purine shortage is a key mechanism underlying the tumor-suppressive effects of DRP-104 (i.e., DON), a glutamine antagonist, across several cancer cell models. This finding aligns with recent studies highlighting the critical roles of purine supply and its disruption in lung cancer models treated with DRP-104^12^, suggesting that this purine shortage-centered mechanism of DRP-104 may be broadly applicable to many cancer types. Most importantly, our work illustrates cellular processes that enable cancer cells to adapt (i.e., develop resistance) to purine shortage and uncovers their vulnerabilities under such a stress. We reveal that cancer cells utilize microtubule-mediated assembly of purinosome-resembling complexes to sustain their purine supply during purine shortage, thereby identifying microtubules as a therapeutic vulnerability in tumor cells under such stress. Additionally, we establish MTAP-dependent purine salvage as a critical source of purine supply in tumor cells, including PCa, lung cancer and glioma cell lines, both under purine-rich conditions and, more notably, during purine shortage (e.g., following DRP-104 treatment). Our findings highlight a novel, protective role of this pathway in tumor cells stressed by DRP-104 treatment. This suggests that targeting the purine salvage pathway could enhance the effects of purine blockade-based therapies. We further demonstrate that purine shortage stress is linked to elevated levels of DNA damage and activation of the cytosolic double-stranded DNA-sensing cGAS-STING pathway, suggesting compromised immune evasion capacity in tumor cells and offering a potential explanation for the DRP-104-induced alterations in the tumor immune microenvironment. Together, these findings inform the mechanisms of DRP-104’s action, the processes underlying tumor cell adaptation/resistance to purine shortage, and future efforts to devise efficacious purine blockade-based therapeutic strategies as described below.

First, while DRP-104 (i.e., DON) is expected to broadly inhibit multiple glutamine-related metabolic enzymes and pathways, its potency against these targets varies, as indicated by the wide range of the Ki values reported in previous studies ^23–32^. Additionally, quantifying the extent to which inhibition of each pathway contributes to tumor-suppressive effects is a challenging task, which can be further compounded by variables such as inhibition kinetics (i.e., doses & timepoints), the strength and roles of specific metabolic pathways in tumor cells, and potential cellular compensation mechanisms. Despite these limitations, the results presented here provide multiple independent lines of evidence to support that purine biosynthesis is among the most consequential processes mediating tumor cell-autonomous suppressive effects of DRP-104 across various cancer types, including PCa, lung cancer and glioma cells. Together with recent findings from the *KEAP1* mutant lung cancer models^12^, these results suggest that purine shortage is an essential mediator of DRP-104’s effects on tumor cells from diverse tissues of origin. We speculate two non-mutually exclusive underlying mechanisms to explain this observation: (i) purine biosynthesis may be the pathway most sensitive to DON (i.e., DRP-104), and/or (ii) purine biosynthesis may hold particular importance for tumor cell propagation compared to other DON-targeted pathways. Further studies are warranted to assess the effects of purine blockade in different cancer types. Additionally, elucidating the interplays between the disruption of purine biosynthesis and glutamine-related metabolic pathways affected by DRP-104 treatment will further enhance our understanding of tumor cell’s adaptation and inform the development of combination therapies.

Second, while the important roles of de novo purines biosynthesis in cancer cells has been well documented^73,74^, recent findings suggest that tumor cells also recycle purines from microenvironment, including dietary sources, to meet their constant need of these essential metabolites^63^. Our findings uncover two new cellular processes that tumor cells employ to ensure sufficient purine supply under purine shortage stress: microtubule-mediated formation of purinosomes - macromolecular complexes composing of multiple de novo purine biosynthesis enzymes that enhance purine biosynthesis ^35,36,39^ - and MTAP-dependent purine salvage. The involvement of purinosomes in tumor cells’ adaptive response to purine biosynthesis inhibition is particularly interesting for two reasons. First, it raises the possibility that these multienzyme complexes may also contribute to tumor progression beyond experimentally induced purine shortage, as highly purine-dependent tumor cells could utilize such complexes to efficiently produce purines to fuel their rapid proliferation. Second, the assembly and proper spatial distribution of purinosomes depend on microtubules and heat shock proteins (e.g., Hsp90 and Hsp70)^38,55,56^. Notably, both microtubules and heat shock proteins are established therapeutic targets in cancer, raising the question of whether drugs targeting these components, such as Docetaxel and Hsp90 inhibitors, might exert part of their anti-tumor effects by impairing purine biosynthesis efficiency, in addition to their known impacts on cell division and oncoprotein functionality. Supported by the effects of Docetaxel described in this study, we suggest that further investigations into this question are warranted, as the outcomes could have direct implications for stratifying cancer patient for specific treatments and understanding mechanisms of resistance to these therapies. For instance, tumors with high purine demand may be more susceptible to these therapies, and reprogramming of purine biosynthesis machinery could potentially underlie resistance to these treatments.

Genetic alterations of MTAP, mostly in the form of homozygous co-deletion with its neighboring tumor suppressor gene *CDKN2A*, are common events in a variety of human cancers ^75,76^. These alterations have been shown to promote tumor progression and aggressiveness, while also suggesting the potential to exploit the resultant MTAP deficiency for treatments ^42,45,47,77–79^. Intriguingly, *MTAP* alterations occur only in a small subset of PCa ^75,76^, supporting the speculation that MTAP may be pro-tumorigenic in prostate cancers^61^. Several previous studies have explored MTAP inhibition for suppressing multiple cancer types, including lung, colon, prostate, and head and neck cancers, by leveraging MTDIA, a potent and specific transition state analogue inhibitor of MTAP ^58–61^. Results from this study suggest that MTAP-mediated purine salvage can serve as an important source of purine supply in prostate and other *MTAP*-intact cancer cells, particularly under conditions of purine shortage. While the roles of MTAP in methionine salvage and the polyamine pathway in PCa pathogenesis remain to be fully addressed, we speculate that MTAP may be pro-tumorigenic in certain contexts through its roles in purine salvage. This aligns with the idea of targeting the MTAP pathway for treating PCa ^61^. Additionally, similar observations regarding MTAP’s role as a purine supply source were made in models of PCa and gliomas. This further supports the idea of exploiting purine blockade to treat cancers with homozygous *MTAP* deletion (i.e., approximately 50% of glioblastomas)^45–47^. Notably, MTAP functions as a purine supply source in cancer types with vastly different frequencies of homozygous *MTAP* deletion; glioblastoma exhibits the highest frequency, while PCa is among the lowest ^75,76^. It is possible that compared to PCa, glioblastoma cells more readily adapt to the loss of purine salvage and/or derive greater benefits from the immunosuppressive microenvironment that MTAP loss may create ^42,64,80^.

Finally, in addition to directly suppressing the propagation of tumor cells, DRP-104 exerts anti-cancer immunity in multiple tumor models in immunocompetent hosts ^10–12^. Previous studies have suggested that MTAP loss contributes to immunosuppressive tumor microenvironment in certain solid tumors ^42,64^. Our findings from in vitro and in vivo tumor models suggest that combination treatment - adding MTDIA to DRP-104 - likely induces equally strong, if not stronger, anti-tumor immunity compared to DRP-104 monotherapy. Most importantly, multiple lines of independent evidence from in vitro and in vivo models support that purine shortage is accompanied by elevated levels of DNA damages and activation of the cytosolic double-stranded DNA-sensing cGAS-STING pathway. Although the precise mechanisms linking these cellular processes remain to be fully elucidated, they offer a potential explanation for the effects of DRP-104 on the tumor immune microenvironment ^10–13^. Mechanisms aside, the potential impact of purine shortage on cancer cells’ DNA damage levels and immune evasion capability has significant therapeutic implications. For example, incorporating DRP-104 into genotoxic treatment regimens could enhance therapeutic efficacy while potentially reducing genotoxicity in normal tissues. Additionally, since the *STING1* gene is often epigenetically silenced in cancers, a process reversible by DNA methyltransferase (DNMT) inhibitors^71^, it is plausible combining DRP-104 with DNMT inhibitors could potentiate anti-tumor effects.

In summary, while the detailed mechanisms underlying cancer cells’ resilience and vulnerability to therapeutically induced purine shortage remain to be further characterized in diverse cellular contexts and in vivo models, we propose that further studies exploring the purine blockade approach in additional prostate cancer models and in other cancer types - including those that frequently harbor *MTAP* loss, such as glioblastoma and lung cancers - are warranted.

## Author Contributions

**Conception and design:** J. Yu, Y He, J Huang

**Methodology development:** J Yu, C Jin, C Pirozzi, X Gao, Y He

**Data acquisition:** J Yu, C Jin, C Su, D Moon, M Sun, X Jiang, N Tserentsoodol, C Pirozzi, X Gao, Y He

**Data analysis and interpretation:** J Yu, C Jin, C Su, D Moon, X Gao, Y He, J Huang

**Writing, reviewing, and revision of the manuscript:** J Yu, C Jin, C Su, D Moon, M Sun, H Zhang, X Jiang, N Tserentsoodol, M Bowie, C Pirozzi, DJ George, R Wild, X Gao, D Ashley, Y He, J Huang

**Administrative, technical, or material support:** J Yu, C Jin, C Su, D Moon, M Sun, H Zhang, X Jiang, F Zhang, N Tserentsoodol, M Bowie, C Pirozzi, DJ George, R Wild, X Gao, D Ashley, Y He, J Huang

**Study supervision:** Y. He, D Ashley, J Huang

## Acknowledgements

We thank Duke Cancer Center Isolation Facility for the support of experiments involving animal models. We thank Dr. Yasheng Gao and the Duke Light Microscopy Core Facility for assisting with imaging experiments, Drs. Darrell Bigner and Vidyalakshmi Chandramohan (Duke University) for sharing reagents, and Dr. Zhiguang Huang for his help with IHC assays. The work was supported by grants from the National Institutes of Health (5R01-CA260726) and the Prostate Cancer Foundation to J.H. We also acknowledge funding support from The Uncle Kory Foundation to Y.H. and NIH/NCI grant R00 CA237618, the USDA 3092-51000-064-05 and Cancer Prevention and Research Institute of Texas (PR210056, CPRIT) Scholar award in Cancer Prevention and Research to X.G.

## Competing Interests

J.H is a consultant for or owns shares in the following companies: Artera, Kingmed Diagnostics, MoreHealth, OptraScan, York Biotechnology, Chimigen Bio, Pfizer and Sisu Pharma.

## Notes

### Summary of Updates

Clinical relevant data added in Supplementary Fig. 3

