## Supplemental figures for "Resilience and vulnerabilities of tumor cells under purine shortage stress"

**Supplementary Figures**

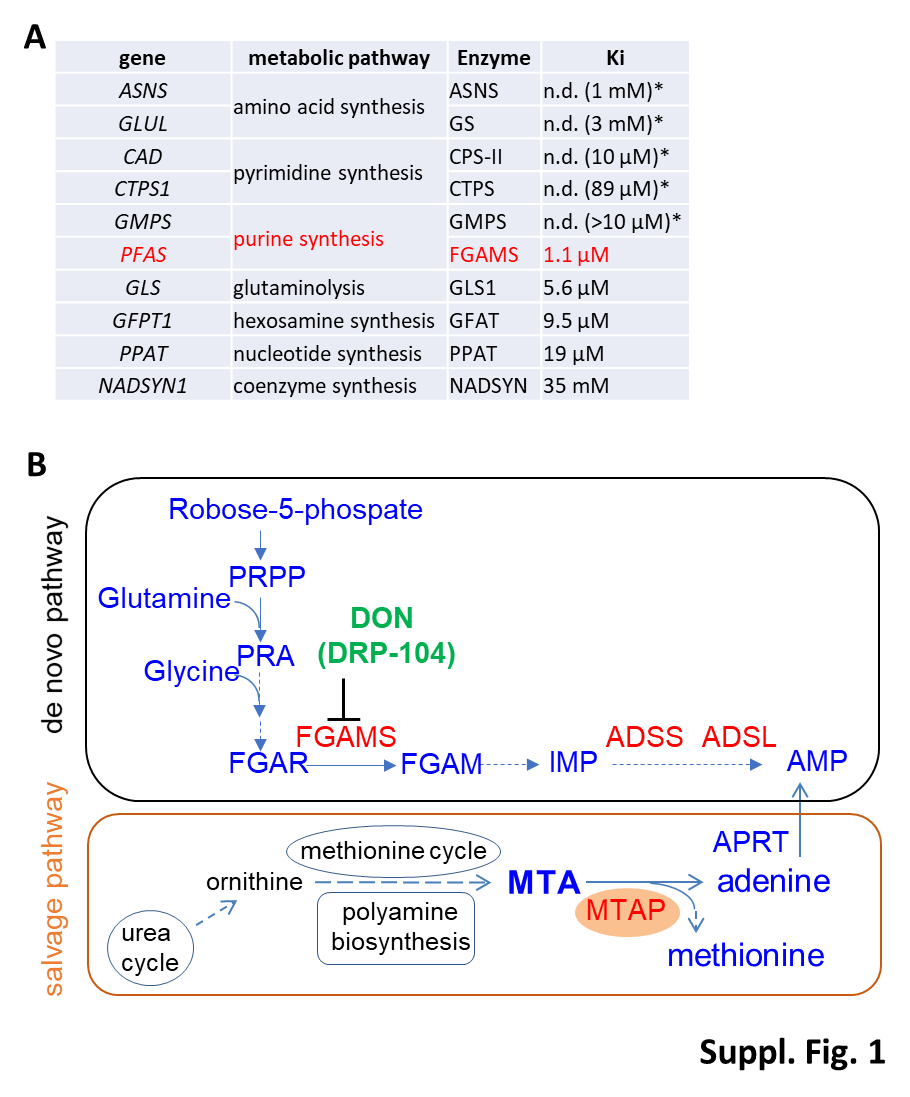

**Supplementary Fig. 1. DON inhibits FGAMS, a de novo purine biosynthesis enzyme, with the lowest Ki. (A)** DON can inhibit 10 enzymes in seven glutamine-utilizing metabolism pathways with variable Ki (*Ki not determined (n.d.); and the concentration reported to be inhibitory was noted). **(B)** Purines can be synthesized in human cells via the de novo pathway (Top) and the salvage pathway (bottom) (metabolites were highlighted in blue and enzymes in red).

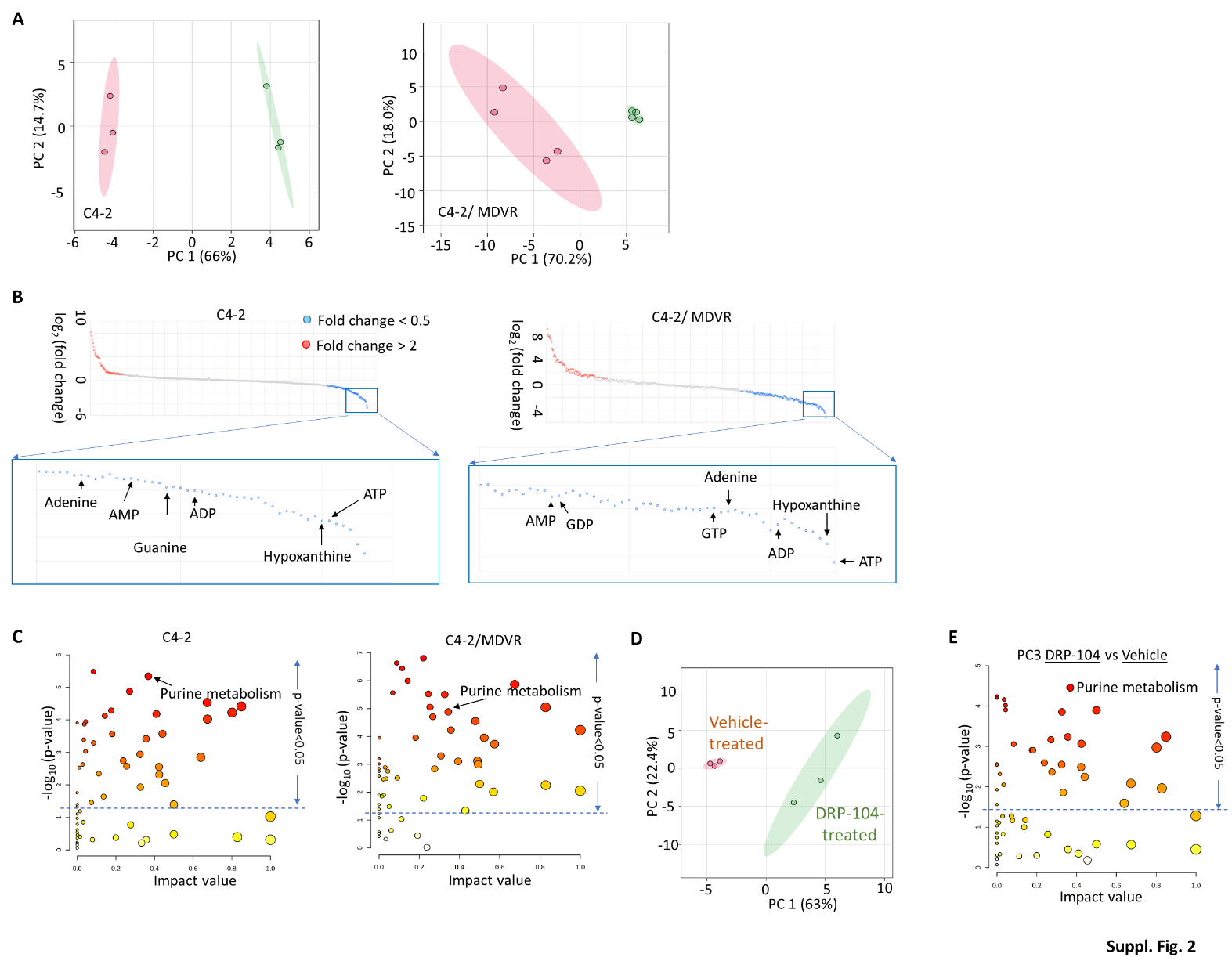

**Supplementary Fig. 2. DRP-104 affects purine metabolism and purine supply. (A-C)** Human CRPC cell line C4-2 or C4-2/MDVR treated with DRP-104 were subjected to metabolic profiling. Principle component analysis distinguished DRP-104-treated cells from vehicle control-treated cells (A). Analysis of the altered metabolites in DRP-104-treated cells vs. the control-treated cells showed that purines were among those that were most strongly affected metabolites (B); and the purine metabolism pathway was among the most affected metabolic pathways (C). (**D-E**) Similar metabolic profiling was performed in PC3 cells and principal component analysis (D) and pathway analysis (E) were shown.

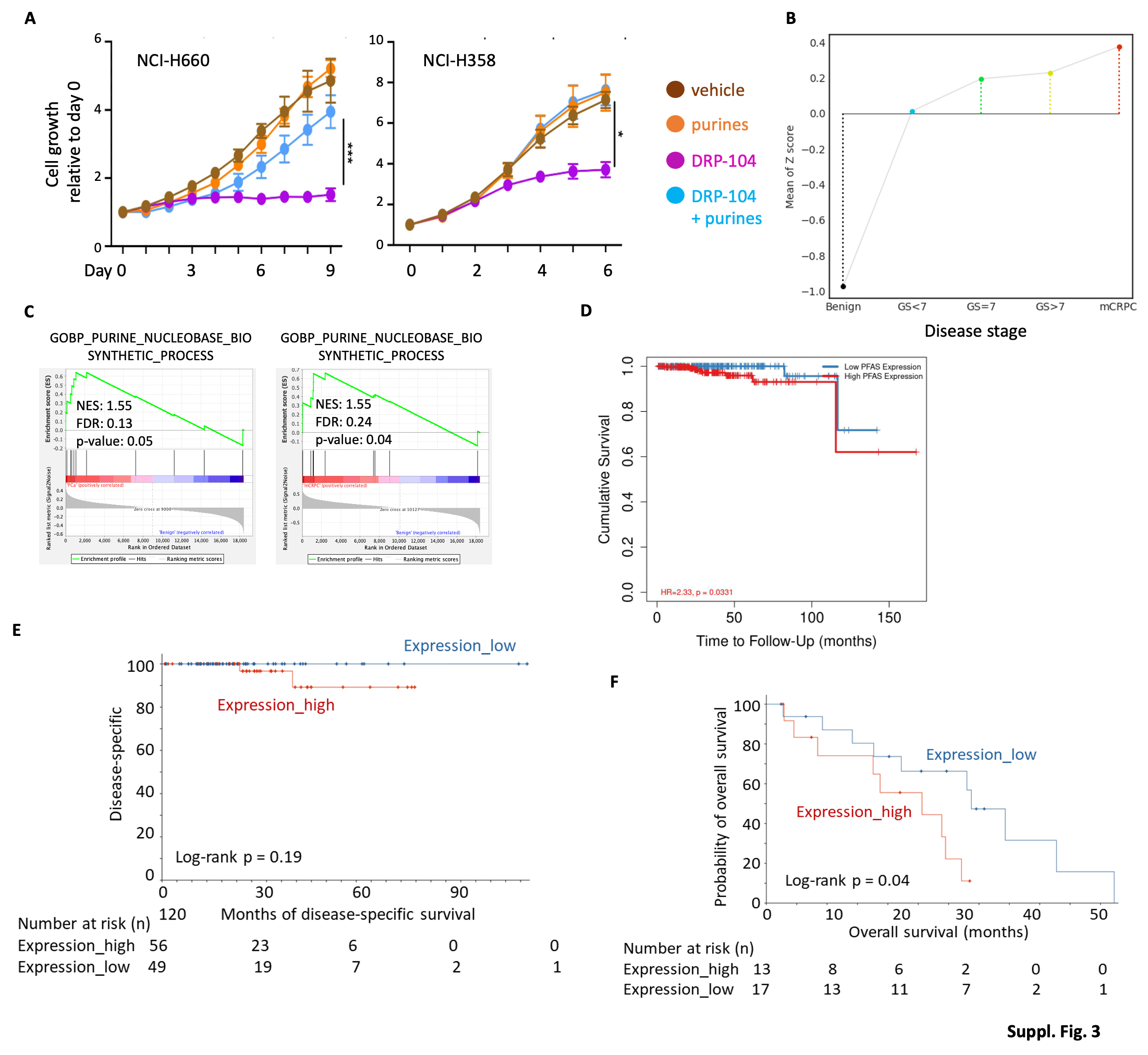

**Supplementary Fig. 3. Gene expression and functional assays implicate purine biosynthesis as a driver of prostate cancer pathogenesis . (A)** Tumor cells were treated with indicated agents and the relative cell proliferation was monitored by IncuCyte. DRP-104 (2 µM) and exogenous purines used included adenosine plus guanosine (each at concentration ranging from 30-100 µM). Data are depicted as mean ± SD. **(B)** A lineplot (generated by http://www.thepcta.org/) shows the mean of Z score for the expression of six key purine biosynthesis enzyme genes (*PPAT*, *GART*, *PFAS*, *PAICS*, *ADSL and ATIC*) in various stages of prostate cancer. One-way ANOVA test between subsets p<0.001; Ranksums-test between primary vs benign p<0.001. **(C)** Gene expression data from (left) primary prostate cancer and (right) mCRPC (from reference ^#^48), in comparison to those from benign tissues, were used for GSEA (GO-Biological Pathway). **(D)** Overall survival curve of PFAS was calculated by the Kaplan–Meier method using TIMER2.0 tools (<http://timer.cistrome.org>) with PFAS median expression as the threshold. **(E)** Disease-specific survival curve of the set of six genes encoding core enzymes of the purine biosynthesis machinery was calculated by the Kaplan–Meier method using cBioPortal tools (<http://www.cbioportal.org/>) in TCGA-PRAD datasets with z-score (relative to normal tissues) >|+/-1.0| as a threshold. Expression_high, at least 1 gene z-score>1.0 without gene z-score<-1.0; Expression_low, at least 1 gene z-score<-1.0 without gene_z-score>1.0. **(F)** Overall survival curve of the set of six genes encoding core enzymes of the purine biosynthesis machinery was calculated by the Kaplan–Meier method using cBioPortal tools (<http://www.cbioportal.org/>) in SU2C dream team datasets with z-score (relative to all samples) >|+/-0.2| as a threshold. Expression_high, at least 1 gene z-score>0.2 without gene z-score<-0.2; Expression_low, at least 1 gene z-score<-0.2 without gene_z-score>0.2. *p<0.05, ***p<0.001.

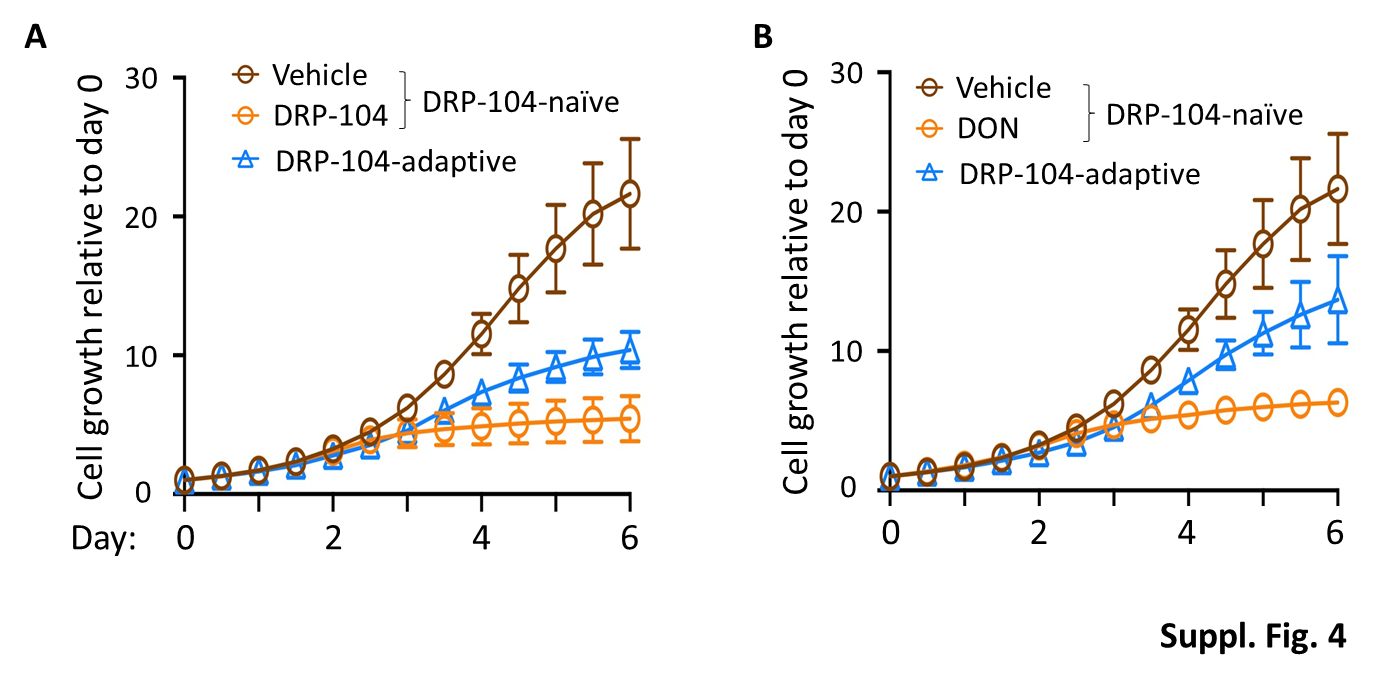

**Supplementary Fig. 4. prostate cancer cells develop adaptation to purine shortage inducer, such as DRP-104.** Cancer cells were treated with DRP-104 (generally in the range of 2-4 µM) to obtain populations that ultimately developed adaptation as defined by their continuous propagation in the presence of this prodrug. In general it takes 3-4 weeks to obtain such “adaptive” cells. Pending on cell lines, the propagation rate of such adaptive cell lines can be identical to (such as C4-2) or slower than (such as PC3) their DRP-104-naïve counterparts. **(A)** The propagation of DRP-104-naïve PC3 cells that were treated with vehicle or with DRP-104 (2 µM), and DRP-104-adaptive PC3 cells that have been maintained in media containing DRP-104 (2 µM). **(B)** The propagation of DRP-104-naïve PC3 cells that were treated with vehicle or with DON (4 µM), and the propagation of DRP-104-adaptive PC3 cells treated with DON (4 µM) (in DRP-104-free media). Data are depicted as mean ± SD.

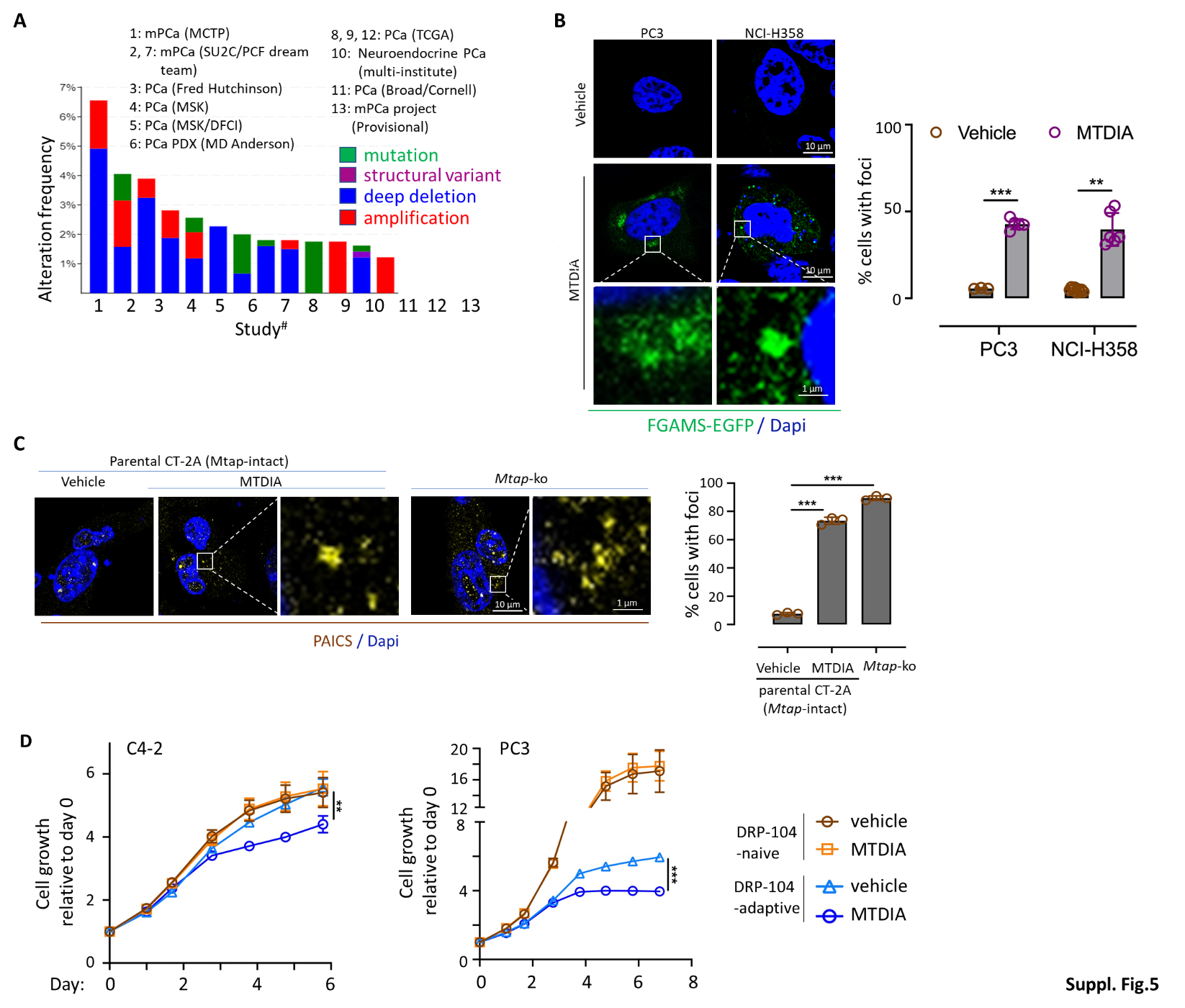

**Supplementary Fig. 5. The MTAP-dependent purine salvage is a source of purine supply for cancer cells. (A)** Oncoprint shows alterations of MTAP in prostate cancer samples (results obtained from cBioportal; only datasets with >1% alteration frequency were shown). Color codes denote types of *MTAP*’s genetic alterations. **(B)** Cancer cell lines (PC3 and NCI-H358) were transfected with exogenous FGAMS-EGFP expressing plasmid and subjected to DRP-104 treatment (4 µM, day 0), and the presence of purinosome foci were determined (day 2). Representative images and quantifications were shown. **(C)** *Mtap*-intact CT-2A cells treated with vehicle control or with MTDIA (5 µM) and *Mtap*-ko CT-2A cells were used for anti-PAICS immunofluorescent staining to detect the presence of purinosomes. Representative images and quantifications were shown. **(D)** Cancer cell lines, including treatment-naïve and their DRP104-adaptive lines, were treated with MTDIA (6 µM) and the relative cell propagation was determined by IncuCyte. Data are depicted as mean ± SEM. **p<0.01, ***p<0.001.

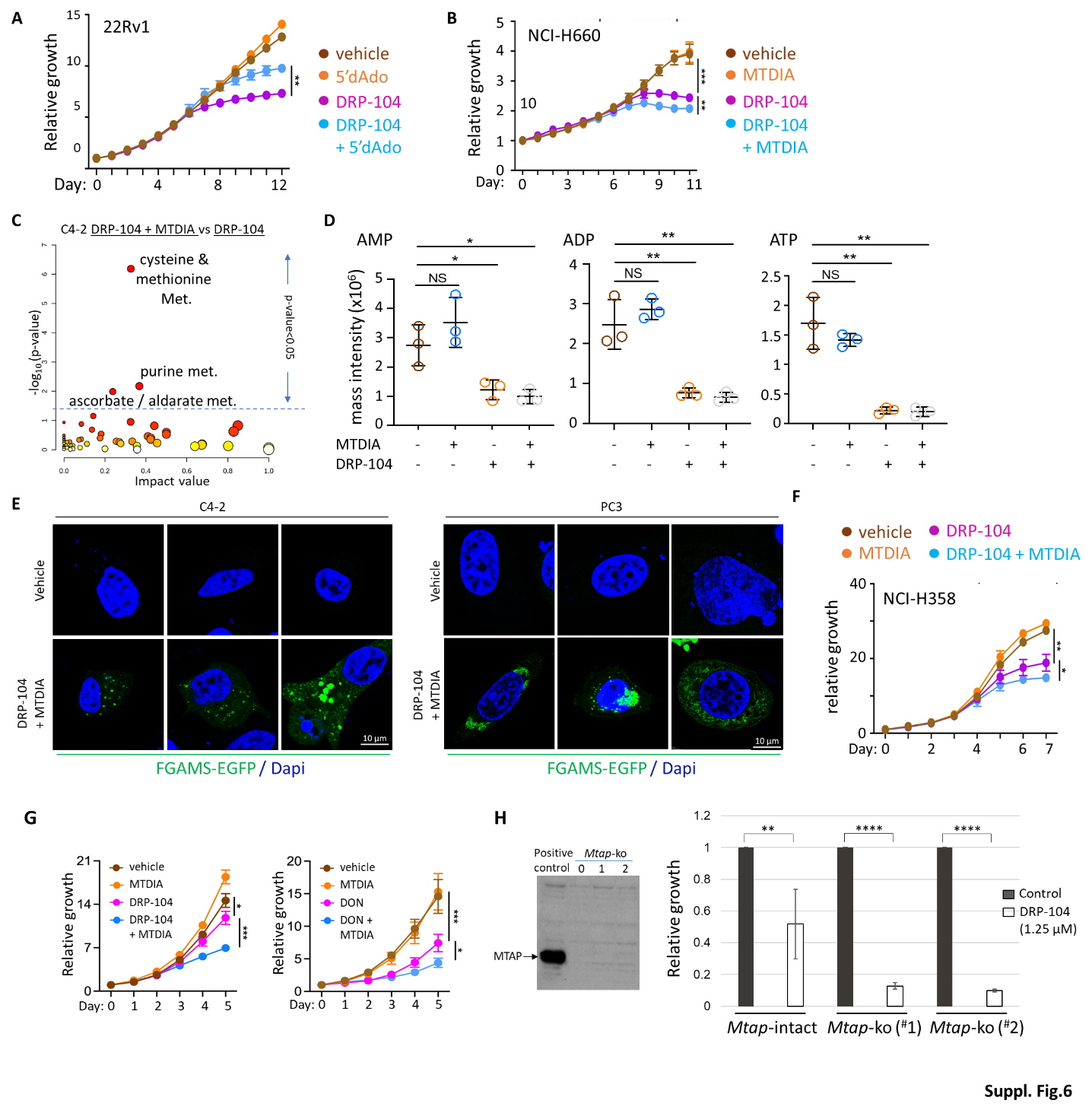

**Supplementary Fig. 6. Inhibiting MTAP sensitizes tumor cells to DRP-104 treatments. (A-B)** Tumor cells were treated with indicated agents and the relative cell proliferation was monitored by IncuCyte. 2 µM of DRP-104 was used for all cell lines; concentrations of 5’dAdo used varied from 10-60 µM, yielding consistent results. Data are depicted as mean ± SD. **(C-D)** C4-2 cells were treated with vehicle control, MTDIA (6 µM) or DRP-104 (5 µM) for two days and metabolite profiling was performed, and the relative abundance of AMP, ADP and ATP was shown. **(E)** CRPC cell lines (C4-2 and PC3) were transfected with exogenous FGAMS-EGFP expressing plasmid and subjected to treatment with DRP-104 plus MTDIA (4 µM and 3 µM respectively, day 0), and the purinosome foci were detected (day 2). Representative images were shown. **(F)** NCI-H358 cells were treated with indicated agents and the relative cell proliferation was monitored by IncuCyte. 2 µM of DRP-104 and 3 or 5 µM of MTDIA were used to obtain similar results. Data are depicted as mean ± SD. **(G)** GL261 cell line was treated with indicated agents and cell proliferation was monitored by IncuCyte (3 µM, 5 µM, and 3 µM was used for DRP-104, DON, and MTDIA, respectively). Data are depicted as mean ± SEM. **(H)** The parental (*Mtap*-intact) GL261 (Redflu-tagged) cell line and its *Mtap*-null derivative lines (*Mtap* loss was confirmed by anti-Mtap immunoblot as shown at the left) were treated with DRP-104 (1.25 µM) and their growth was determined (relative cell growth on day 6 was shown; standard deviations were denoted). *p<0.05, **p<0.01, ***p<0.001, ****p<0.0001.

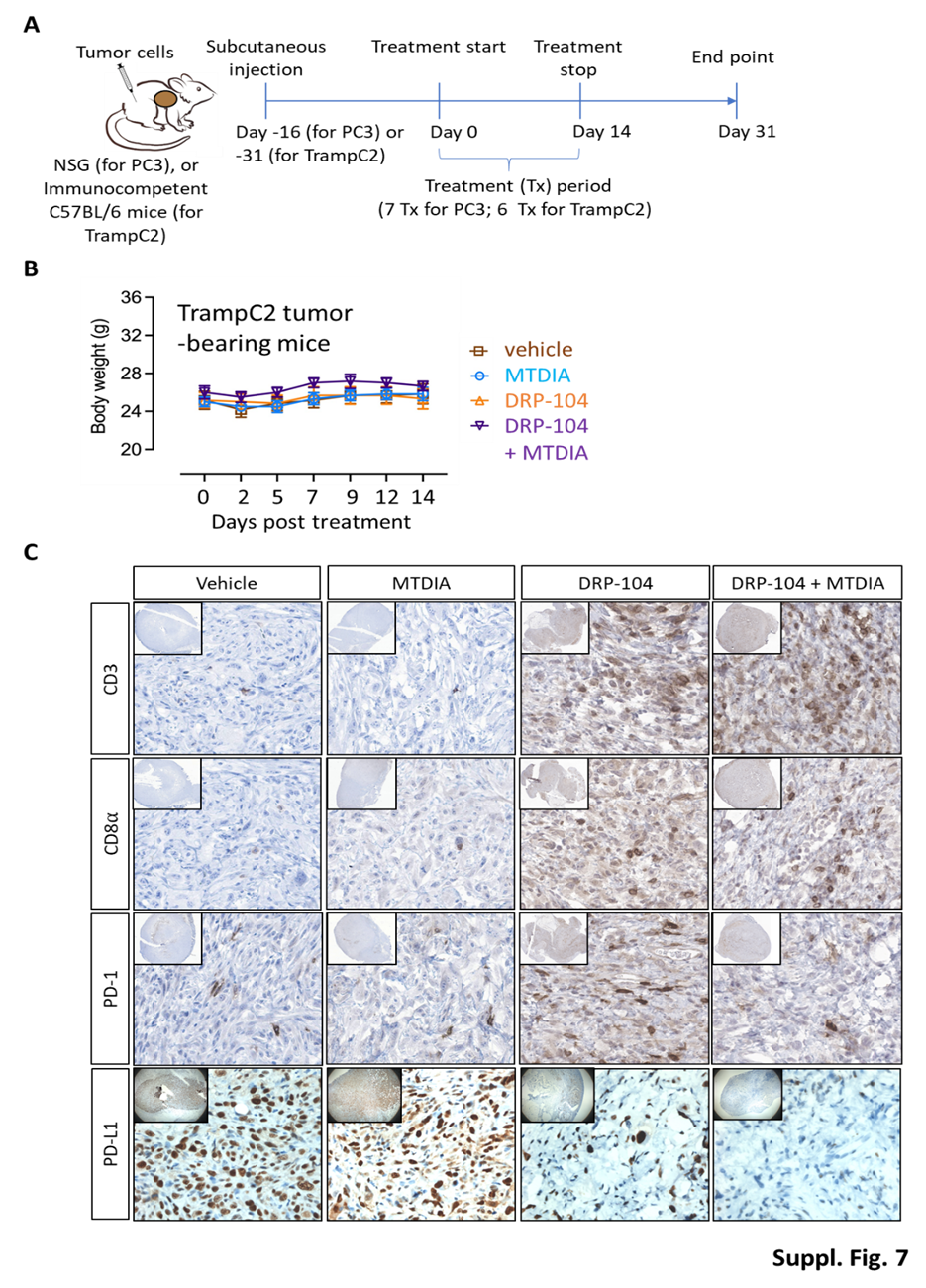

**Supplementary Fig. 7. Treating tumor-bearing mice with DRP-104 in combination with MTDIA does not affect animal’s body weight but alters the tumor’s immune microenvironment. (A)** Treatment regimens. **(B)** The body weights of mice (bearing TrampC2 tumors) during the course of treatment were determined and plotted (n=6 per arm). Data are depicted as mean ± SEM. **(C)** TrampC2 tumor tissues harvested from immune competent host mice after the end of two week’s treatments were used for IHC detection of indicated marker proteins. Representative images were shown.

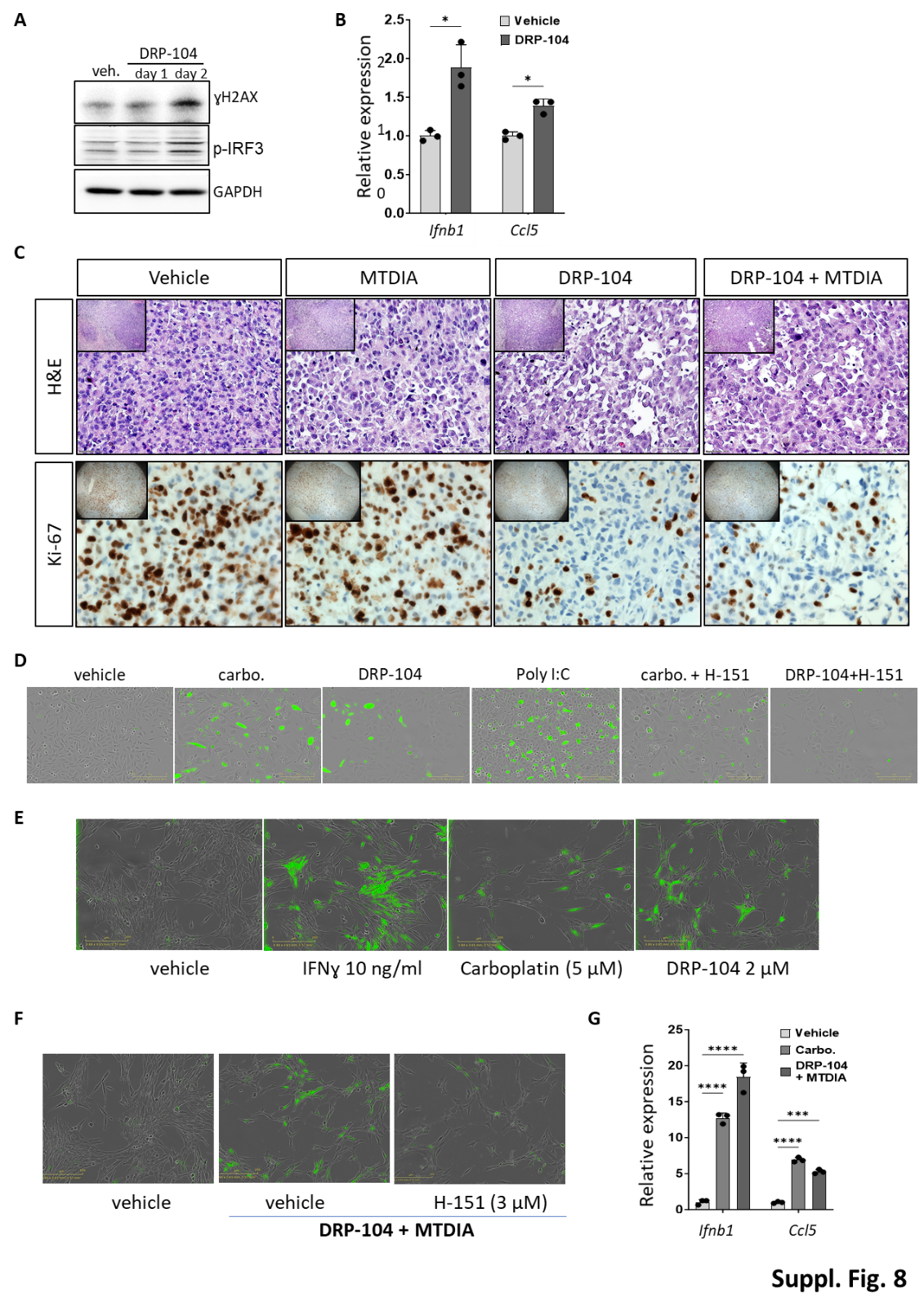

**Supplementary Fig. 8. DRP-104 simulates DNA damage and the cGAS-STING pathway. (A-B)** TrampC2 cells was treated with DRP-104 (2 µM) and cells were harvested for **(A)** immunoblots and **(B)** RT-qPCR for gene expression assays. **(C)** Representative H&E staining and anti-Ki67 IHC images from PC3-derived xenografts tumors after they were treated with indicated agents for two weeks and harvested at their endpoints. Note the presence of tumor cells regardless the treatment group and the reduced abundance of Ki67+ cells in the DRP-104-treated and DRP-104+MTDIA-treated groups at the endpoints (when the tumors were not actively treated). **(D)** ISRE reporter-tagged PC3 cells were treated with indicated agents and representative images showing the reporter activation were shown (poly I:C, a known stimulator of the ISRE activity, was included as a positive control). **(E-F)** ISRE reporter-tagged CT-2A cells were treated with indicated agents and representative images showing the activation of the reporter signal were shown (IFNɣ was included as a positive control). **(G)** Parental CT-2A cells were treated with indicated agents and the expression of marker genes in the cGAS-STING pathway, *Ifnb1* and *Ccl5*, was determined. carbo: carboplatin. *p<0.05, ****p<0.0001.
