## Supplementary materials for "Resilience and vulnerabilities of tumor cells under purine shortage stress"

**Reagents and kits for Yu et. al.**

| **Product name** | **Company** | **Catalog Number** | **Applications** |
| --- | --- | --- | --- |
| DMEM | Thermo Fisher | Cat# 10564011 | Cell Culture |
| RPMI 1640 | Thermo Fisher | Cat# 11875093 | Cell Culture |
| Advanced DMEM/F-12 | Thermo Fisher | Cat# 12634010 | Cell Culture |
| Fetal bovine serum | Thermo Fisher | Cat# 26140079 | Cell Culture |
| Recombinant Human EGF | Peprotech | Cat# AF-100-15 | Cell Culture |
| B-27™ Supplement | Thermo Fisher | Cat# 17504044 | Cell Culture |
| Recombinant Human FGF | peprotech | Cat# 100-18B | Cell Culture |
| L-glutamine | Thermo Fisher | Cat# 11965092 | Cell Culture |
| Human Recombinant Insulin | Sigma‒Aldrich | Cat# 91077C | Cell Culture |
| Dihydrotestosterone | Millipore Sigma | Cat# D073 | Cell Culture |
| Penicillin‒streptomycin | Thermo Fisher | Cat# 15140122 | Cell Culture |
| Phosphate-Buffered Saline | Thermo Fisher | Cat# 10010049 | Cell Culture |
| Trypsin-EDTA (0.25%), phenol red | Thermo Fisher | Cat# 25200114 | Cell Culture |
| StemPro™ Accutase | Thermo Fisher | Cat# A1110501 | Cell Culture |
| Dimethyl sulfoxide (DMSO) | Millipore Sigma | Cat# D2650 | Cell Culture |
| Mycoplasma Detection Kit | InvivoGen | Cat# rep-mysnc-50 | Cell Culture |
| Lipofectamine 3000 | InvitroGen | Cat# L3000015 | Transfection |
| µ-Slide 4 Well | iBidi | Cat# 80426 | IF |
| Polyformaldehyde | Santa Cruz Biotechnology | Cat# sc-281692 | IF |
| Triton X-100 | Bio-Rad | Cat# 1610407 | IF |
| DAPI | MilliporeSigma | Cat# D9542 | IF |
| ProLong™ Gold Antifade Mountant | Thermo Fisher | Cat# P36930 | IF |
| T4 Polynucleotide Kinase | New England Biolabs | Cat# M0201S | Molecular clone |
| Esp3I (BsmBI) | Thermo Fisher | Cat# ER0451 | Molecular clone |
| Macherey-Nagel™ NucleoSpin™ kit | Thermo Fisher | Cat# 11982382 | Molecular clone |
| Quick Ligation™ Kit | New England Biolabs | Cat# M2200S | Molecular clone |
| NEB® Stable Competent E. coli | New England Biolabs | Cat# C3040H | Transformation |
| Q5® Hot Start High-Fidelity 2X Master Mix | New England Biolabs | Cat# M0494S | PCR Amplification |
| QIAprep Spin Miniprep Kit | QIAGEN | Cat# 27106 | Plasmid Extraction |
| TransIT-X2® | Mirus Bio | Cat# MIR 6004 | Virus Construction |
| PEG 8000 | Thomas Scientific | Cat# C752W61 | Virus Concentration |
| 5M NaCl | Sigma‒Aldrich | Cat# S6546 | Virus Concentration |
| Polybrene | Millipore Sigma | Cat# TR1003 | Transduction |
| Puromycin | Millipore Sigma | Cat# P8833 | Selection |
| Quick-DNA Miniprep Plus Kit | ZYMO RESEARCH | Cat# D4069 | gDNA Extraction |
| RIPA buffer | Sigma‒Aldrich | Cat# R0278 | WB |
| Protease and phosphatase inhibitor cocktail | Thermo Fisher | Cat# 1861281 | WB |
| 4x Laemmli Sample Buffer | Bio-Rad | Cat# 161074 | WB |
| Immun-Blot PVDF membranes | Bio-Rad | Cat# 1620177 | WB |
| Pierce ECL substrate | Thermo Fisher | Cat# 21050 | WB |
| SuperSignal West Femto Maximum Sensitivity Substrate | Thermo Fisher | Cat# 34095 | WB |
| Immun-Blot PVDF Membrane Roll | Bio-Rad | Cat# 1620177 | WB |
| TRI Reagent® | Sigma‒Aldrich | Cat# T9424 | RNA extraction |
| Quick-RNA Miniprep Kit | ZYMO RESEARCH | Cat# R1055 | RNA extraction |
| High-Capacity cDNA Reverse Transcription Kit | Thermo Fisher | Cat# 4368814 | qRT-PCR |
| SYBR Green qPCR Master Mix | YEASEN | Cat# 11184ES08 | qRT-PCR |
| Ultra Hematoxylin | Azer scientific | Cat# SKUES36101 | H&E |
| ULTRA EOSIN-Y | Azer scientific | Cat# SKUES36111 | H&E |
| Citrate Buffer | Millipore Sigma | Cat# C9999 | IHC |
| Peroxidazed 1 | BiocareMedical | Cat# PX968M | IHC |
| Background Punisher | BiocareMedical | Cat# BP974L | IHC |
| DAKO Antibody Diluent | Agilent | Cat# S302283-2 | IHC |
| Mouse HRP | Agilent | K4001 | IHC |
| Rabbit HRP | Agilent | K4003 | IHC |
| DAKO Bluing Buffer | Agilent | Cat# CS70230-2 | IHC |
| DAKO DAB | Agilent | K3468 | IHC |

**Abbreviations:** IF: Immunofluorescence; qRT-PCR: Quantitative Real-Time Polymerase Chain Reaction; WB: Western Blotting; H&E: Hematoxylin and eosin; IHC: Immunohistochemistry.

**Antibodies**

| **Antibodies** | **Company** | **Catalog Number** | **Applications** | **Dilutions** |
| --- | --- | --- | --- | --- |
| PAICS | Proteintech | Cat# 12967-1-AP | IF | 1:200 |
| anti-rabbit Alexa 594 | ThermoFisher | Cat# A-11037 | IF | 1:500 |
| β-Actin | Cell Signaling Technology | Cat# 4967 | WB | 1:1000 |
| GAPDH | Cell Signaling Technology | Cat# 2118 | WB | 1:2000 |
| α-Tubulin | Cell Signaling Technology | Cat# 3873 | WB | 1:1000 |
| Vinculin | Themo Fisher | Cat# 14-9777-82 | WB | 1:1000 |
| AR | Abcam | Cat# ab74272 | WB | 1:1000 |
| PSA | Cell Signaling Technology | Cat# 5365S | WB | 1:1000 |
| γ-H2AX | Cell Signaling Technology | Cat# 9718 | WB | 1:1000 |
| H2AX | Cell Signaling Technology | Cat# 7631 | WB | 1:1000 |
| p-IRF3 (Ser386) | Cell Signaling Technology | Cat# 37829 | WB | 1:1000 |
| Phospho-IRF3 (Ser396) | Themo Fisher | Cat# MA5-14947 | WB | 1:1000 |
| Anti-IRF3 (Ser386) | Abcam | Cat# ab192796 | WB | 1:1000 |
| IRF3 | Cell Signaling Technology | Cat# 11904 | WB | 1:1000 |
| MTAP | ABclonal | Cat# A20907 | WB | 1:1000 |
| GART | Proteintech | Cat# 2A11E2 | WB | 1:7000 |
| Goat Anti-Mouse IgG (H+L)-HRP Conjugate | Bio-Rad | Cat# 1721011 | WB | 1:7000-10000 |
| Goat Anti-Rabbit IgG (H + L)-HRP Conjugate | Bio-Rad | Cat# 1706515 | WB | 1:7000-10000 |
| Phospho-IRF3 (Ser396) | Themo Fisher | Cat# MA5-14947 | IHC | 1:50 |
| γ-H2AX | Cell Signaling Technology | Cat# 9718 | IHC | 1:50 |
| CD3 | BOSTER | Cat# PB9093 | IHC | 1:100 |
| CD8⍺ | BOSTER | Cat# A02236-1 | IHC | 1:100 |
| PD-1 | BOSTER | Cat# A00178 | IHC | 1:100 |
| PD-L1 | Proteintech | Cat# 17952-1-AP | IHC | 1:500 |

**Abbreviations:** IF: Immunofluorescence; WB: Western Blotting; IHC: Immunohistochemistry.

**Primers for RT-qPCR:**

| **Target Gene** | **Forward** | **Reverse** |
| --- | --- | --- |
| *Actb* | GGTGGGAATGGGTCAGAAGG | GTACATGGCTGGGGTGTTGA |
| *Ifnb1* | GCCTTTGCCATCCAAGAGATGC | ACACTGTCTGCTGGTGGAGTTC |
| *Ccl5* | CCTGCTGCTTTGCCTACCTCTC | ACACACTTGGCGGTTCCTTCGA |
| *ACTB* | ATAGCACAGCCTGGATAGCAACGTAC | CACCTTCTACAATGAGCTGCGTGTG |
| *IFNB1* | TTGTTGAGAACCTCCTGGCT | TGACTATGGTCCAGGCACAG |
| *CCL5* | CCTGCTGCTTTGCCTACATTGC | ACACACTTGGCGGTTCTTTCGG |
